# Mitochondrial DNA damage in substantia nigra pars compacta astrocytes exacerbates dopaminergic neuron loss in a 6-hydroxydopamine mouse model of parkinsonism

**DOI:** 10.1101/2025.08.30.673287

**Authors:** Daniela A. Ayala, Grace M. Hall, Eliana Tijerina, Luisa Lopez, Zaid Khan, Devanshi Paliwal, Anu Anand, Mendell Rimer, Rahul Srinivasan

**Author notes:** Corresponding author at (R. Srinivasan).

## Abstract

Parkinson’s disease (PD) is the fastest growing neurological disorder with no known cure. Our ability to develop disease-modifying treatments that slow down the loss of substantia nigra pars compacta (SNc) dopaminergic (DA) neurons is hindered by a dearth of knowledge on roles for non-neuronal elements such as astrocytes during PD pathogenesis. More specifically, the extent to which mitochondrial DNA (mtDNA) damage in SNc astrocytes contributes to SNc DA neuron loss during PD remains unknown. To address this knowledge gap, we utilized an adeno-associated virus (AAV) called Mito-PstI that expresses the restriction enzyme PstI as an approach to damage mtDNA in SNc astrocytes and assess the effect of astrocytic mtDNA damage on SNc DA neuron function and viability in mice. Mito-PstI-induced mtDNA damage in SNc astrocytes disrupted mitochondrial morphology, caused increased wrapping of SNc astrocytic processes around SNc DA neurons, and abnormally increased dopamine release by SNc DA neuron axonal terminals within the dorsolateral striatum (DLS). In addition, mice injected with Mito-PstI in the SNc showed increased spontaneous and apomorphine-induced rotations contralateral to the side with SNc Mito-PstI injections. In further experiments, we used a parkinsonian mouse model with low dose 6-hydroxydopamine (6-OHDA) injection into the DLS to show that Mito-PstI expression in SNc astrocytes caused a worsening of 6-OHDA-induced spontaneous contralateral rotational behavior, and exacerbated SNc DA neuron loss. These results suggest that mitochondria in SNc astrocytes are not only critical for the function of SNc DA neurons, but are also a new target for developing disease-modifying strategies against PD.

**Graphical Abstract:** 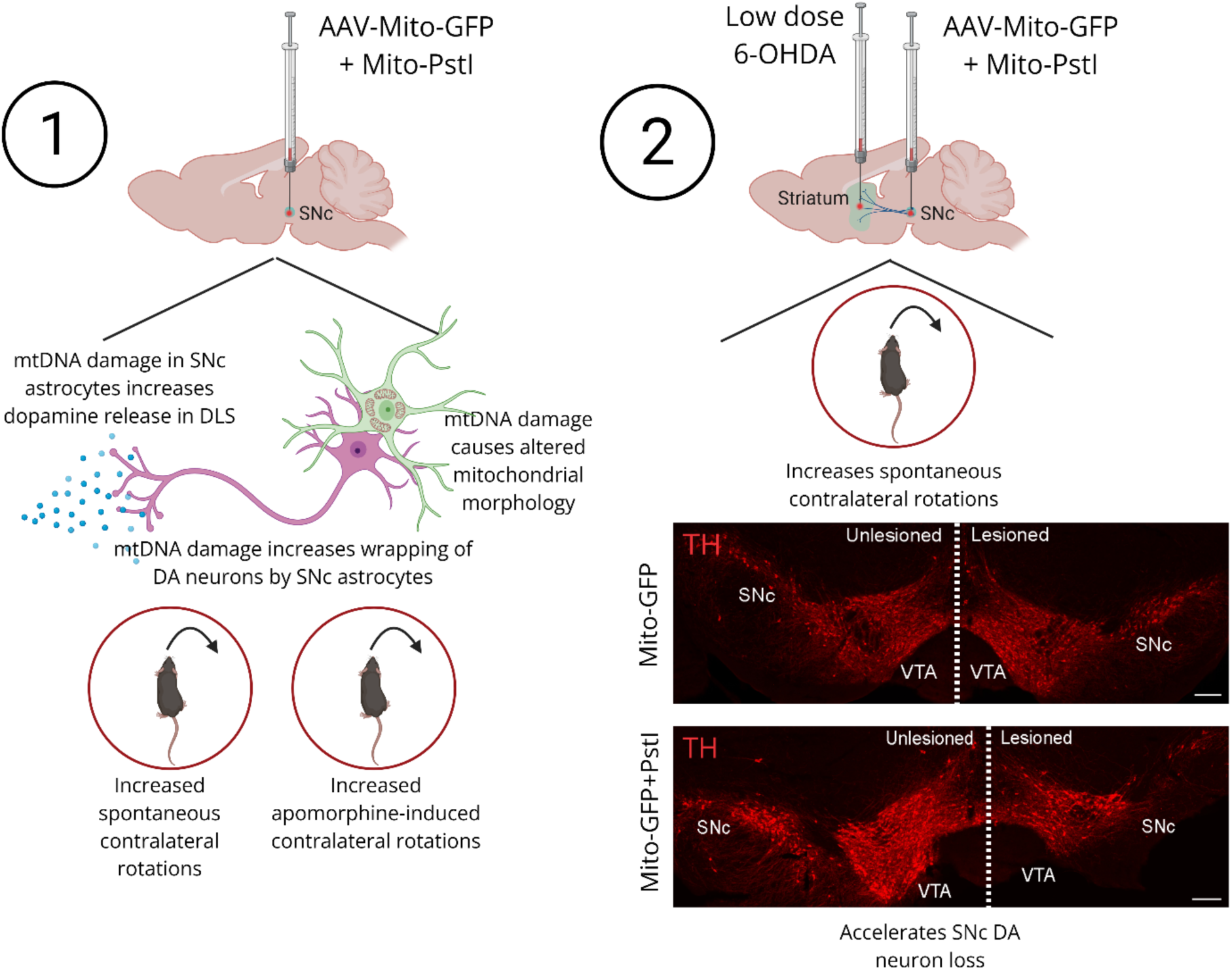

**Main Points:**

- Mito-PstI expression in mouse SNc astrocytes increases dopamine release
- Mito-PstI expression in SNc astrocytes worsens contralateral rotational behavior in mice
- Mito-PstI expression in SNc astrocytes increases DA neuron loss in 6-OHDA mice

## Introduction

Parkinson’s disease (PD) is the fastest growing neurological disorder (Dorsey, Sherer, Okun, & Bloem, 2018; Su et al., 2025) with no known cure. Since the motor dysfunction observed in clinical PD is primarily caused by an accelerated loss of substantia nigra pars compacta (SNc) dopaminergic (DA) neurons, disease modifying strategies for PD generally focus on protecting SNc DA neurons from death, with the objective of slowing down the progression of this devastating neurodegenerative disease (Hirsch, Jenner, & Przedborski, 2013; Lindholm et al., 2016).

One knowledge gap that hinders our ability to develop clinically effective disease-modifying treatments for PD is that we lack a comprehensive understanding of the contribution of non-cell autonomous mechanisms during PD pathogenesis (Pons-Espinal, Blasco-Agell, & Consiglio, 2021). This is important because in contrast to neuron-specific cell autonomous mechanisms that affect individual DA neurons within the SNc, non-cell autonomous mechanisms can cause dysfunction across the entire SNc, thereby leading to rapid DA neurodegeneration (Dawson, 2008; di Domenico et al., 2019; Gleichman & Carmichael, 2020). In this context, astrocytes within the SNc are an important non-neuronal target to consider during PD pathogenesis because these cells not only outnumber neurons (Gonzalez-Perez, Lopez-Virgen, & Quinones-Hinojosa, 2015) but can also become reactive in PD (Booth, Hirst, & Wade-Martins, 2017; Smajic et al., 2022). In addition, multiple studies suggest that astrocytes play a pathogenic role in neurodegeneration via aberrant calcium signaling (Bancroft & Srinivasan, 2021), mutations in PD-related risk genes within astrocytes (Booth et al., 2017), altered processing of α-synuclein (Ozoran & Srinivasan, 2023), as well as abnormal interactions with microglia and the blood brain barrier (Garcia, Zarate, & Srinivasan, 2024).

Although there are a multitude of proposed pathological mechanisms by which astrocytes can accelerate SNc DA neuron loss and the progression of PD (Garcia et al., 2024), the specific role of SNc astrocytic mitochondria in contributing to neurodegeneration is a relatively unexplored, but active area of investigation (Bantle, Hirst, Weihofen, & Shlevkov, 2020; Gollihue & Norris, 2020). Astrocytic mitochondria are particularly relevant to SNc DA neuron loss for multiple reasons: (**i**) The brain alone consumes 20% of the total energy required by the body, and central nervous system (CNS) mitochondria meet a majority of this massive energy requirement for the brain (Cunnane et al., 2020; Panov, Orynbayeva, Vavilin, & Lyakhovich, 2014). (**ii**) Mitochondria in CNS astrocytes are strategically distributed within subcellular regions in the astrocytic territory, endfeet and soma (Ayala et al., 2024; Huntington & Srinivasan, 2021; Zarate, Huntington, Bagher, & Srinivasan, 2023), and since astrocytes generally outnumber neurons in the CNS (Gonzalez-Perez et al., 2015), astrocytic mitochondria are optimally positioned to cater to the high energy demands of CNS neurons, and particularly, SNc DA neurons. (**iii**) Compromised function in astrocytic mitochondria could have profound detrimental effects on neuronal function within the SNc, which is corroborated by evidence showing that SNc astrocytes demonstrate deficiencies in mitochondrial oxidative phosphorylation (OXPHOS) protein complex expression in postmortem samples of patients with clinically diagnosed PD (Chen et al., 2022). (**iv**) Astrocytic mitochondria are highly active organelles, displaying a wide panorama of Ca^2+^ influx events that robustly respond to neurotransmitter agonists (Huntington & Srinivasan, 2021). (**v**) Mutations in PD-related risk genes that are present in astrocytes have been shown to cause mitochondrial dysfunction, which can potentially be detrimental to surrounding neurons (Booth et al., 2017). Taken together, these multiple reports motivate the important question of whether specifically damaging mitochondria in SNc astrocytes could accelerate SNc DA neuron loss during PD.

Based on the rationale presented above, this study directly assesses the role of an important non-neuronal element, *viz.* SNc astrocytic mitochondria, and more specifically, astrocytic mitochondrial DNA (mtDNA) damage in accelerating SNc DA neuron loss in mice. We utilize our newly developed adeno-associated virus (AAV)-based tool (Ayala et al., 2024) to show that mtDNA damage in SNc astrocytes alters their mitochondrial morphology along with alterations in the physical interaction between astrocytes and DA neurons in the SNc. This is accompanied by dramatic increases in acetylcholine (ACh)-evoked dopamine release by SNc DA neuron terminals in the dorsolateral striatum (DLS), a worsening of 6-hydroxydopamine (6-OHDA) induced behavioral deficits in mice, and an exacerbation of SNc DA neuron loss following exposure to 6-OHDA. Together, the results presented here pinpoint SNc astrocytic mitochondria as a translationally relevant non-neuronal disease-modifying target to slow SNc DA neuron loss and the progression of PD.

## Materials and Methods

### Mice

Two to three-month old C57BL/6 wildtype male and female mice (Jackson Laboratories) were used for all experiments. Mice were housed on a 12 h light/dark cycle with *ad libitum* access to food and water. All experiments were performed in accordance with Texas A&M University Institutional Animal Care and Use Committee approved guidelines (IACUC #2022-0252).

### AAV constructs

Mito-GFP (AAV2/5-GfaABC1D-mito7-GFP) and Mito-PstI (AAV2/5-GfaABC1D-mito7-PstI) constructs were generated by Vector Builder (Chicago, IL). AAV-GRABDA (AAV9-hsyn-GRAB_DA2m) was used to express the optogenetic sensor GRABDA in the DLS and was obtained from Addgene (Cat. # 140553). For all AAV injections, the injection volume was ~1 μl AAV, suspended in PBS. Mice were injected with either 2×10^10^ genome copies (gc) of Mito-GFP into the SNc (control mice), or co-injected with 1×10^10^ gc of Mito-GFP + 1×10^10^ gc of Mito-PstI (experimental mice) into the SNc. For imaging of acetylcholine (ACh)-evoked dopamine release in the dorsolateral striatum (DLS) of live brain slices, mice were injected with 1×10^13^ gc of AAV-GRABDA into the DLS, 7 days prior to AAV injections of either the Mito-GFP (control mice) or Mito-GFP + Mito-PstI (experimental mice) into the SNc on the same side as striatal AAV-GRABDA injections.

### Experimental protocols

This study utilizes three separate experimental protocols in mice for assessing the effect of PstI expression on SNc DA neuron physiology, behavior, SNc astrocytic mitochondria, SNc astrocyte-neuron interactions and neurodegeneration. A detailed timeline for each of the protocols is depicted in the figures.

The first protocol assesses the effect of PstI expression in SNc astrocytic mitochondria on ACh-evoked dopamine release from SNc DA neuron terminals in the DLS in mice. For this protocol, mice were injected unilaterally into the DLS with AAV9-hsyn-GRAB_DA2m (called AAV-GRABDA), and 7 days later, the same mice were injected with either Mito-GFP (in control mice) or Mito-GFP + Mito-PstI (in experimental mice) into the SNc, on the same side as DLS injection of AAV-GRABDA. Live imaging of dopamine release in mouse striatal slices was performed 28 days after DLS injection of AAV-GRABDA, using changes in GRABDA fluorescence as a quantitative readout.

The second protocol longitudinally assesses the effect of PstI expression in astrocytic mitochondria on SNc DA neurons of mice without 6-OHDA exposure. For this protocol, spontaneous and apomorphine-induced contralateral rotations were performed on days −7 (baseline), 21, 28, 35, and 42. Day 0 was the time point for injecting either the AAV 2/5 GfaABC1D-mito7-GFP (called Mito-GFP) (control mice) or Mito-GFP + AAV 2/5 GfaABC1D-mito7-PstI (called Mito-PstI) (experimental mice).

The third protocol longitudinally assesses the extent to which Mito-PstI expression in SNc astrocytic mitochondria of mice accelerates 6-OHDA-induced SNc DA neuron loss. For this protocol, spontaneous and apomorphine-induced contralateral rotations were performed on days −7 (baseline), 14, 28, 35, and 42, where day 0 was the time point for unilaterally injecting either the Mito-GFP (control mice) or Mito-GFP + Mito-PstI (experimental mice) into the SNc, and day 21 was the time point for unilateral 6-OHDA injection into the DLS on the same side as AAV injections into the SNc. For protocols 2 and 3, all mice were perfused at the end of the experiment, after final rotational behavior was completed on day 42. For the last two protocols, midbrain sections were collected from perfused mice to assess mitochondrial morphology, astrocytic wrapping of SNc DA neurons and SNc DA neuron loss with tyrosine hydroxylase (TH) staining.

### Stereotaxic Surgeries

Stereotaxic surgeries were performed to inject mice with AAVs into the SNc and DLS as well as 6-OHDA into the DLS (Sigma, St. Louis, MO). For unilateral 6-OHDA lesions in the DLS, 5 mg/ml stock solutions of 6-OHDA (Sigma, St. Louis, MO) were prepared in 0.9% saline with 0.2% ascorbic acid and frozen at −80°C until use. Mice received a unilateral stereotaxic injection of 0.6 μl of stock solution (3 μg of 6-OHDA in total per mouse) into the DLS at a rate of 0.25 μl/min. Stereotaxic surgeries were performed as previously described (Ayala et al., 2024; Huntington & Srinivasan, 2021; Zarate, Garcia, Pandey, & Srinivasan, 2025; Zarate et al., 2023; Zarate et al., 2021). All surgeries were performed on 2 to 3-month-old male and female mice under general anesthesia using isoflurane, with induction at 5% and maintenance at 1-2%, dispensed through a SomnoSuite Low Flow Anesthesia System (Kent Scientific, Torrington, CT). All injections were done using a beveled glass injection pipette, attached to a Stoelting Quintessential Stereotaxic Injector (QSI), at a flow rate of 0.3 μL/min. SNc coordinates were −3.0 mm posterior to bregma, −1.5 mm lateral to bregma, and −4.2 mm ventral to pial surface. DLS coordinates were +0.9 mm anterior to bregma, +1.8 mm lateral to bregma, and +2.5 mm ventral to the pial surface. The injection pipette was left in place for 5-10 minutes after injection, then gradually withdrawn, and surgical wounds were closed with tissue adhesive (Amazon).

### Imaging of ACh-evoked changes of GRABDA fluorescence in live mouse DLS brain slices

ACh-evoked changes in GRABDA fluorescence within the DLS was used as a readout to quantify dopamine release from DLS axonal terminals of SNc DA neurons in mice following injections of control Mito-GFP and experimental Mito-PstI AAVs into the SNc. For live slice imaging, mice were sacrificed 28 days after injection of AAV-GRABDA into the DLS. Mice were decapitated using isoflurane and brains were rapidly extracted. 300 μm-thick coronal slices of the DLS were obtained using a vibratome (Ted Pella Inc., D.S.K Microslicer ZERO 1 N). The slicing solution consisted of (in mM): 194 sucrose, 30 NaCl, 4.5 KCl, 1.2 NaH_2_PO_4_, 26 NaHCO_3_, 10 D-glucose, and 1 MgCl_2_ and saturated with 95% O_2_ and 5% CO_2_ (pH 7.2). Slices were incubated in a solution of 25% slicing solution and 75% ACSF recording solution at 34°C for ~20 minutes, then maintained at room temperature throughout the experiment. All recordings were performed in an artificial cerebrospinal fluid (ACSF) recording solution consisting of (in mM): 126 NaCl, 2.5 KCl, 1.24 NaH_2_PO_4_, 26 NaHCO_3_, 10 D-glucose, 2.4 CaCl_2_, and 1.3 MgCl_2_ saturated with 95% O_2_ and 5% CO_2_ (pH 7.4). Live-slice imaging was performed using an Olympus FV3000 upright laser-scanning confocal microscope equipped with a 40x water immersion objective (N.A. 0.45) with a 3x optical zoom. 488 nm LED-based excitation wavelength was used to visualize GRABDA fluorescence in the DLS. During all live imaging sessions, confocal parameters such as LED power, voltage, gain, offset and aperture diameter remained constant. The sampling rate was 1 frame per second (FPS) for 600 seconds. ACh (Sigma, cat#A2661-25G) was bath perfused using a peristaltic pump (Harvard Apparatus). Bath perfusion timing was set prior to and maintained constant at each imaging session. To assess changes in dopamine release in the DLS, spontaneous activity was recorded for the first 180 seconds, followed by the bath application of 300 μM ACh for 420 seconds. All recording solutions contained 500 nM of the acetylcholinesterase inhibitor, donepezil (Sigma, cat#D6821) to prevent rapid degradation of bath applied ACh.

### Acquisition and quantification of spontaneous and apomorphine-induced rotational behavior in mice

Apomorphine-induced rotational behavior was performed as previously described (Zarate et al., 2025; Zarate et al., 2021). Mice were individually placed in 5-gallon circular buckets (33 cm in diameter, one mouse per bucket) and spontaneous rotational behavior was recorded for 10 min. Following acquisition of spontaneous rotational behavior, the same mice were injected intraperitoneally (i.p) with apomorphine (0.5 mg/kg in 0.9% saline; Sigma, cat #1041008) and contralateral rotational behavior was video recorded for 10 min immediately after apomorphine injections. All videos were recorded with a ceiling mounted camcorder. The total number of contralateral rotations occurring during the 10 min. period was calculated using the Ethovision XT program (Noldus). All detection settings on Ethovision were maintained constant across cohorts of mice.

### Immunostaining

Forty-two days after the AAV injections in protocols 1 and 2 as described in the methods section above, mice were deeply anesthetized with isoflurane, then transcardially perfused with PBS, immediately followed by 10% formalin (VWR, cat# 100496-506). After extraction, brains were preserved in 10% formalin at 4°C for 48 hours, then dehydrated in 30% sucrose in PBS (Sigma, cat# S7903) for 48 h. Forty micrometer coronal midbrain sections were obtained using a Epredia, CryoStar NX50 cryostat and stored in 0.01% sodium azide in PBS (Sigma, cat # S2002). To maintain precise rostro-caudal orientation, midbrain sections were sequentially collected in 96-well plates. Midbrain sections were washed in 1X PBS, prior to being blocked and permeabilized in 10% normal goat serum (NGS, Abcam, ab7481) and 0.5% Triton X-100 (Sigma, cat# 9002-93-1) rocking gently for 45 min at room temperature. Sections were washed twice with PBS before overnight incubation at 4°C in primary antibodies. For astrocyte wrapping, process density analysis and quantification of SNc DA neurons loss using TH labeling, chicken anti-TH (1:1000, Abcam cat# ab76442) and rabbit anti-S100B (1:1000, Abclonal cat# a19108) primary antibodies were used. Sections were washed twice the following day with 1X PBS followed by incubation in secondary antibodies goat anti-chicken Alexa Fluor 594 (1:2000, Abcam cat# ab150176) and goat anti-rabbit ATTO 647N (1:2000, Rockland cat# 611-156-122S).

### Imaging of immunostained midbrain sections

To quantify SNc DA neuron loss using TH labeling, fixed tissue sections were imaged using an Olympus VS120 Virtual Slide Scanning System equipped with a UPlanSApo 20X air objective. 561 nm LED-based excitation with an exposure time of 400 ms per section was used for obtaining images from multiple midbrain sections per mouse. For astrocyte wrapping and process density analysis using S100B labeling, 40 μm thick midbrain sections were imaged using an Olympus FV3000 laser-scanning confocal microscope. Images were obtained using a 60x oil immersion objective (N.A. 0.8) with 3x optical zoom to focus on individual SNc DA neurons along with astrocytic mitochondria and S100B-labeled processes in their immediate vicinity. LED-based excitation wavelengths (in nm) at 488 (endogenous Mito-GFP labeled SNc astrocytic mitochondria), 561 (TH labeled SNc DA neurons), and 647 (S100B labeled astrocytic processes) were used for acquiring images. All confocal images consist of ~20 to 30 optical sections with a 0.5 μm step size at 3x digital zoom. Confocal parameters such as excitation intensity, HV, gain and offset were maintained constant for each condition.

### Quantification of mitochondrial morphology in SNc astrocytes

Three parameters of mitochondrial morphology, *viz.* mitochondria count, mitochondrial area, and mitochondria size were quantified per FOV from images that were obtained using a 60x oil immersion objective (N.A. 0.8) with 3x optical zoom to focus on individual SNc DA neurons along with astrocytic mitochondria surrounding the SNc DA neuron soma. AAV-expressed Mito-GFP labeling in SNc astrocytic mitochondria enabled clear visualization of the mitochondrial structure in SNc astrocytes surrounding SNc DA neuronal soma within the FOV. All parameters for mitochondrial morphology were quantified using ImageJ (ver.1.54f). For quantification, a max intensity z-stack was created for each SNc FOV. A rectangular region of interest was then made encompassing the entire FOV. The FOV was smoothed and manually thresholded to create ROIs of all mitochondria particles within the FOV by using the analyze particle function in ImageJ. Individual mitochondrial particle ROIs within each FOV were used to obtain the average mitochondrial count, mitochondrial network area, and mitochondrial particle size per FOV.

### Quantification of astrocytic S100B wrapping around SNc DA neurons

To quantify astrocyte wrapping of SNc DA neurons, 2-dimensional projections of z-stacks of TH and S100B labeled midbrain sections were generated using the ImageJ maximum intensity function. A region of interest (ROI) for SNc DA neuron soma was defined by manually tracing the TH-labeled neuronal somas and applying to the S100B maximum intensity projection. In this case, all S100B signals outside the neuronal ROI area were deleted to focus on S100B labeled SNc astrocyte process on only the SNc DA neuron soma. Using this method, S100B labeling within the SNc DA neuron soma ROI was manually thresholded, and a second ROI was created to demarcate astrocyte processes that closely overlay SNc DA neuron somatic ROIs. Using a ratio of the S100B ROI area over SNc DA neuron soma area, the astrocytic S100B wrapping percent on each SNc DA neuron soma was calculated by dividing the S100B labeled process area by the soma area and multiplying by 100 to acquire a wrapping index, expressed in percent. In addition, astrocyte process density for the whole field of view (FOV) was determined by dividing the thresholded astrocyte process area by the total FOV area and multiplying by 100 to obtain a percent density of S100B-labeled astrocytic processes in each FOV.

### Quantification of ACh-evoked dopamine release in the DLS

To quantify ACh-evoked dopamine release, confocal XY time-series movies were drift-corrected along the x-y axis in ImageJ. An ROI encompassing the entire FOV was used to obtain mean gray values of GRABDA fluorescence intensities across the entire recording time of 600 s, and 600 s time traces of mean GRABDA fluorescence in intensities were generated for each FOV. Using the integrate tool from the OriginLab gadget menu, area under the curve (AUC) of GRABDA fluorescence intensity from time traces were obtained during the 420 s period of ACh bath application for each FOV, which was from 180 to 600 s during each recording. For consistency, the 420 s window with bath applied ACh for measuring mean GRABDA fluorescence AUC values was maintained constant across all live brain slice FOVs and mice.

### Quantification of DA neuron loss in the SNc

SNc DA neuron loss was quantified using TH immunolabeling as a marker for SNc DA neurons. Three independent readouts were used for quantification of SNc DA neuron loss: (**i**) TH+ SNc DA cell number, (**ii**) TH+ SNc DA neuron area, and (**iii**) Intensity of TH fluorescence in immunolabeled SNc DA neurons. To quantify each of these three readouts, Z-stack projections of TH labeled midbrain sections for each mouse were generated using ImageJ sum slices function. Using projected Z-stacks of midbrain sections, the number of SNc TH+ neurons within 6-OHDA lesioned and unlesioned and / or AAV-injected and uninjected sides were manually counted in 4 evenly spaced midbrain sections per mouse, and the neuron number ratio between 6-OHDA lesioned and unlesioned and / or AAV-injected and uninjected SNc was obtained for each midbrain section, and multiple mice per condition. For TH area and fluorescence intensity quantification, Z-projected images from each midbrain section were manually thresholded to isolate TH+ cell bodies and processes. Thresholded images were used to define ROIs on the 6-OHDA lesioned and unlesioned and / or AAV-injected and uninjected sides of the SNc. Area and integrated fluorescence intensity was obtained for the thresholded ROI area on the 6-OHDA lesioned and unlesioned and / or AAV-injected and uninjected SNc of each midbrain section. Data from 4 sections per mouse were obtained, and ratios between the 6-OHDA lesioned and unlesioned and / or AAV-injected and uninjected SNc was obtained per section for area and integrated intensity.

### Statistical analyses

All statistical analyses were performed using OriginLab. For comparison of two conditions, the Shapiro-Wilk test was first used to test data for normality. Normally distributed datasets were compared using a two-sample t-test, while non-normal datasets were tested using the Mann-Whittney test. For statistical testing of rotational behavior, a three-way analysis of variance (three-way ANOVA) was used to examine interactions between three factors that were treatment (Mito-GFP versus Mito-GFP + PstI), sex (males and females) and time (days of testing). A *p*-value that was less than 0.05 was considered as statistically significant. Unless otherwise indicated, data are expressed as mean ± SEM. Results, figures and figure legends describe exact *p* values, sample sizes and specific statistical tests used for each experiment. For 3-way ANOVA, F statistics and the *p* value for main interaction effect are reported in the results. Because we did not observe any sex differences, we combined male and female mice for all experimental readouts in this study.

## Results

### Mito-PstI alters the morphology of mitochondria in SNc astrocytes

We previously developed an AAV-based tool to specifically damage mtDNA in astrocytes *in vivo* in a targeted, brain-region specific manner (Ayala et al., 2024). Our novel AAV called Mito-PstI, expresses the restriction enzyme PstI under control of the astrocyte-specific GfaABC1D promoter. AAV-expressed PstI is directed to the astrocytic mitochondrial matrix by a mitochondria-specific mito7 targeting signal sequence (Mito), where this restriction enzyme cuts mtDNA at PstI recognition sites in mice, thereby damaging mtDNA only in astrocytes within pre-specified Mito-PstI injected brain regions *in vivo* (Ayala et al., 2024). We used this novel AAV-based method to determine the extent to which SNc astrocytic mtDNA damage alters the morphology of mitochondria in mouse SNc astrocytes.

The SNc of mice was stereotaxically injected with AAVs expressing either Mito-GFP alone (control mice) or Mito-GFP + Mito-PstI (experimental mice) and 3 weeks later, midbrain sections obtained from AAV-injected mice were immunostained for TH to visualize GFP-labeled astrocytic mitochondria surrounding TH+ SNc DA neurons (Fig. 1A). Using high-resolution confocal imaging, we found that both Mito-GFP and Mito-GFP + Mito-PstI injected mice displayed SNc astrocytic mitochondria in close proximity to TH+ SNc DA neuron soma (Fig. 1B and C). When compared to Mito-GFP control mice, we found that co-expression of Mito-GFP + Mito-PstI in the SNc caused a significant ~2-fold increase in the number of SNc astrocytic mitochondrial particles (Average number of mitochondrial particles per FOV for Mito-GFP = 186.15 ± 13.48 and Mito-GFP + Mito-PstI = 363.93 ± 26.23; *p* < 0.0001, Mann-Whitney test), and a corresponding ~2-fold increase in total SNc astrocytic mitochondrial area (Average area of mitochondria per FOV for Mito-GFP = 95.95 ± 8.31 μm^2^ and Mito-GFP + Mito-PstI = 203.12 ± 17.23 μm^2^; *p* < 0.0001, Mann-Whitney test). However, there was no significant change in the size of SNc astrocytic mitochondrial particles (Average size of mitochondrial particles per FOV for Mito-GFP = 0.472 ± 0.016 μm^2^ and Mito-GFP + Mito-PstI = 0.506 ± 0.023 μm^2^; *p* = 0.336, Mann-Whitney test) (Fig. 1D–F).

**Figure 1.**
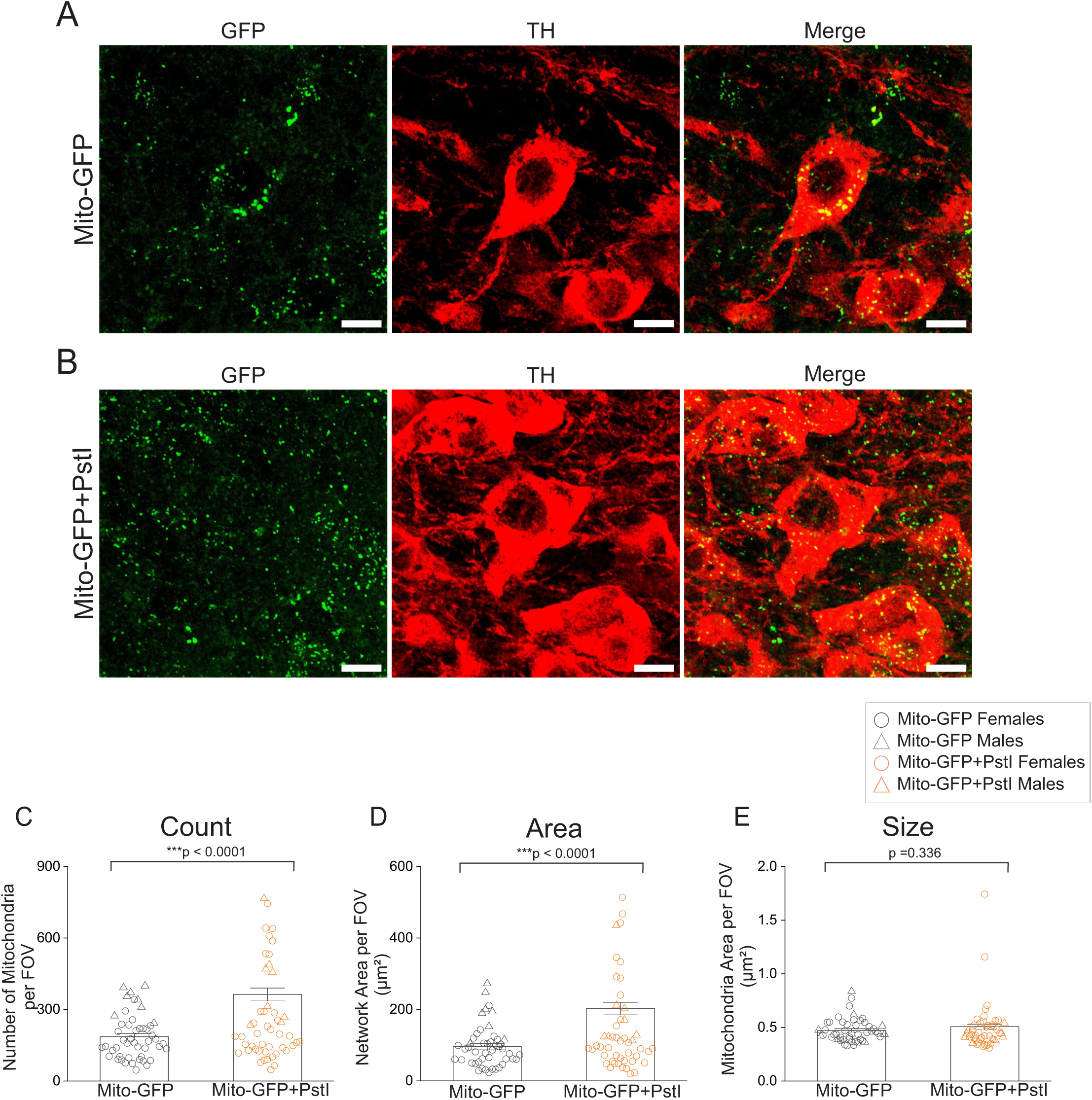
Mito-PstI expression alters mitochondrial morphology in SNc astrocytes. (**A**) Representative z-stack projections of confocal images from immunostained SNc midbrain sections showing Mito-GFP (green), TH (red) in the SNc and merged image of Mito-GFP and TH. Scale bar = 10 μm. (**B**) Representative confocal images of immunostained SNc midbrain sections showing Mito-GFP + Mito-PstI (green), TH (red) and merged image of Mito-GFP + Mito-PstI and TH. Scale bar = 10 μm. (**C**) Bar graph comparing GFP-labeled mitochondrial particle count between Mito-GFP (control) and Mito-GFP + Mito-PstI (experimental) in the SNc of AAV injected mice (**D**) Bar graph comparing GFP-labeled mitochondrial area per FOV between Mito-GFP (control) and Mito-GFP + Mito-PstI (experimental) in the SNc of AAV injected mice (**E**) Bar graph comparing average GFP-labeled mitochondrial particle size per FOV between Mito-GFP (control) and Mito-GFP + Mito-PstI (experimental) in the SNc of AAV injected mice. n = 48 FOVs from 4 mice for Mito-GFP and n = 72 FOVs from 6 mice for Mito-GFP + Mito-PstI. Individual FOV data points from male and female mice for each condition are shown as circles for females and triangles for males. Data expressed as mean ± SEM, *p*-values are based on a Mann-Whitney test.

### Mito-PstI increases the wrapping of astrocyte processes around TH+ DA neurons in the SNc

Having observed specific changes in the morphology of SNc astrocytic mitochondria following Mito-PstI expression (Fig. 1), we asked if astrocytic mtDNA damage via Mito-PstI expression in SNc astrocytes can alter the physical interaction of astrocytic processes in relation to TH+ SNc DA neuron soma. To do this, the mouse SNc was stereotaxically injected with AAVs expressing either Mito-GFP (control mice) or Mito-GFP + Mito-PstI (experimental mice). Three weeks later, coronal midbrain sections obtained from AAV-injected mice were co-immunostained for SNc astrocyte-specific S100B and TH (Fig. 2A and B). Using our previously published method (Bancroft, De La Mora, Pandey, Zarate, & Srinivasan, 2022), we quantified the extent of wrapping of S100B labeled SNc astrocyte processes around DA neurons in AAV-injected mice. When compared to control mice expressing Mito-GFP, experimental mice co-expressing Mito-GFP + Mito-PstI in the SNc showed a significant ~16 % increase in the wrapping of S100B containing astrocytic processes around TH+ SNc DA neurons (Fig. 2C) (% wrapping of S100B+ astrocyte processes around TH+ DA neuron soma for Mito-GFP = 43.57 ± 1.71 and for Mito-GFP + Mito-PstI = 51.10 ± 1.29%; *p* < 0.0001, Mann Whitney test). Based on our finding that wrapping of S100B+ astrocyte processes around SNc DA neuron soma was significantly increased (Fig. 2C), we asked if Mito-PstI also caused a general increase in the percent density of SNc astrocyte processes. Indeed, we found that Mito-GFP + Mito-PstI-co-injected mice showed a significant ~17 % increase in S100B+ process density per FOV when compared to control Mito-GFP-injected mice (Fig. 2D) (% S100B+ astrocyte process density per FOV for Mito-GFP = 34.97 ± 0.918 and for Mito-GFP + PstI = 42.09 ± 0.8877; p < 0.0001, Mann-Whitney test). To further assess if the observed morphological changes in SNc astrocyte processes (Fig. 2C and D) was associated with an increase in astrocyte reactivity, we compared average S100B intensity in the SNc between control Mito-GFP injected and experimental Mito-GFP + Mito-PstI mice. When compared to Mito-GFP mice, Mito-GFP + Mito-PstI injected mice showed a significant ~15 % reduction in S100B intensity (Fig. 2E) (Average S100B intensity per FOV for Mito-GFP = 428.20 ± 10.54 A.U. and for Mito-GFP + PstI = 364.43 ± 9.90 A.U.; p < 0.0001, Mann-Whitney test). Together, these data suggest that Mito-PstI-induced changes in the morphology of SNc astrocytic mitochondria are associated with morphological changes in the interaction between astrocyte processes and SNc DA neurons, but with no discernable increase in SNc astrocyte reactivity.

**Figure 2.**
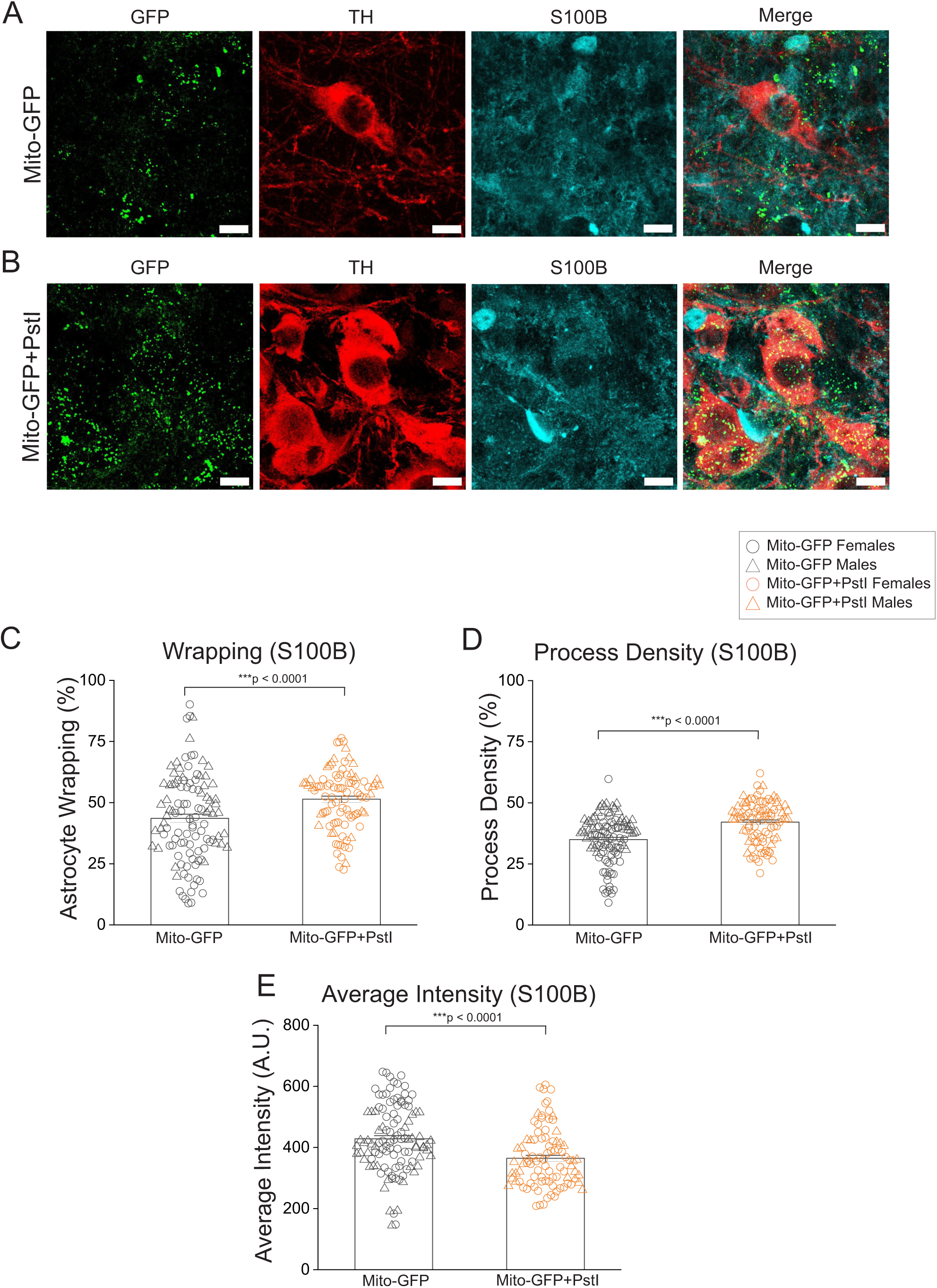
Mito-PstI expression in the SNc increases wrapping of astrocyte processes around SNc DA neuron somata. (**A**) Representative z-stack projections of confocal images from immunostained SNc midbrain sections showing Mito-GFP (green), TH (red) and S100B (cyan) and merged images of Mito-GFP, TH and S100B. Scale bar = 10 μm. (**B**) Representative z-stack projections of confocal images of immunostained SNc midbrain sections showing Mito-GFP + Mito-PstI (green), TH (red) and S100B (cyan) and merged image of Mito-GFP + Mito-PstI, TH and S100B. Scale bar = 10 μm. (**C**) Bar graph showing percent S100B wrapping around SNc DA neurons per FOV (**D**) Bar graph showing S100B process density in the SNc surrounding TH+ SNc DA neurons per FOV (**E**) Bar graph showing average S100B intensity in astrocytes surrounding TH+ SNc DA neurons per FOV. n = 109 FOVs from 12 mice for Mito-GFP and n = 92 FOVs from 6 mice for Mito-GFP + Mito-PstI. Data expressed as mean ± SEM, *p*-values are based on a Mann-Whitney test. Individual FOV data points from male and female mice for each condition are shown as circles for females and triangles for males.

### Mito-PstI expression in the SNc increases ACh-evoked dopamine release in the DLS

We rationalized that because Mito-PstI-induced mtDNA damage in SNc astrocytes causes a significant increase in the wrapping of astrocytic processes around SNc DA neurons (Fig. 2C), this could potentially increase astrocytic clearance of dopamine released from the somatodendritic compartment of SNc DA neurons, causing dysregulated axonal dopamine release in the DLS via a homeostatic imbalance in autoregulatory mechanisms governing axonal dopamine release (Ford, 2014). Based on this rationale, we sought to directly assess changes in ACh-evoked dopamine release from DA axonal terminals within the DLS of live mouse brain slices following expression of either control Mito-GFP or experimental Mito-GFP + Mito-PstI in SNc astrocytes.

The DLS of mice was first injected with an AAV expressing GRABDA, which is an optogenetic sensor for dopamine (Sun et al., 2018). One week later, AAVs expressing either Mito-GFP (control mice) or Mito-GFP + Mito-PstI (experimental mice) were injected into the SNc (Fig. 3A). Three weeks following SNc AAV injections, ACh-evoked changes in GRABDA fluorescence were used to quantify dopamine release in the DLS of live striatal mouse brain slices (Fig. 3A). We found that when compared to control SNc Mito-GFP expressing mice, bath application of 300 μm ACh in striatal slices of experimental SNc Mito-GFP + Mito-PstI co-expressing mice showed a dramatic ~40 % greater increase in GRABDA fluorescence as measured with AUC (Fig. 3B–D, Supplementary Movies 1 and 2) (ACh-evoked changes in GRABDA AUC: Mito-GFP = 1761.46 ± 213.92 A.U. x sec and Mito-GFP + PstI = 2942.43 ± 403.077 A.U. x sec; *p* = 0.015, Mann Whitney test). These data show that the total amount of releasable dopamine within DA axons in the DLS of mice is abnormally increased following Mito-PstI expression-induced mtDNA damage in SNc astrocytes.

**Figure 3.**
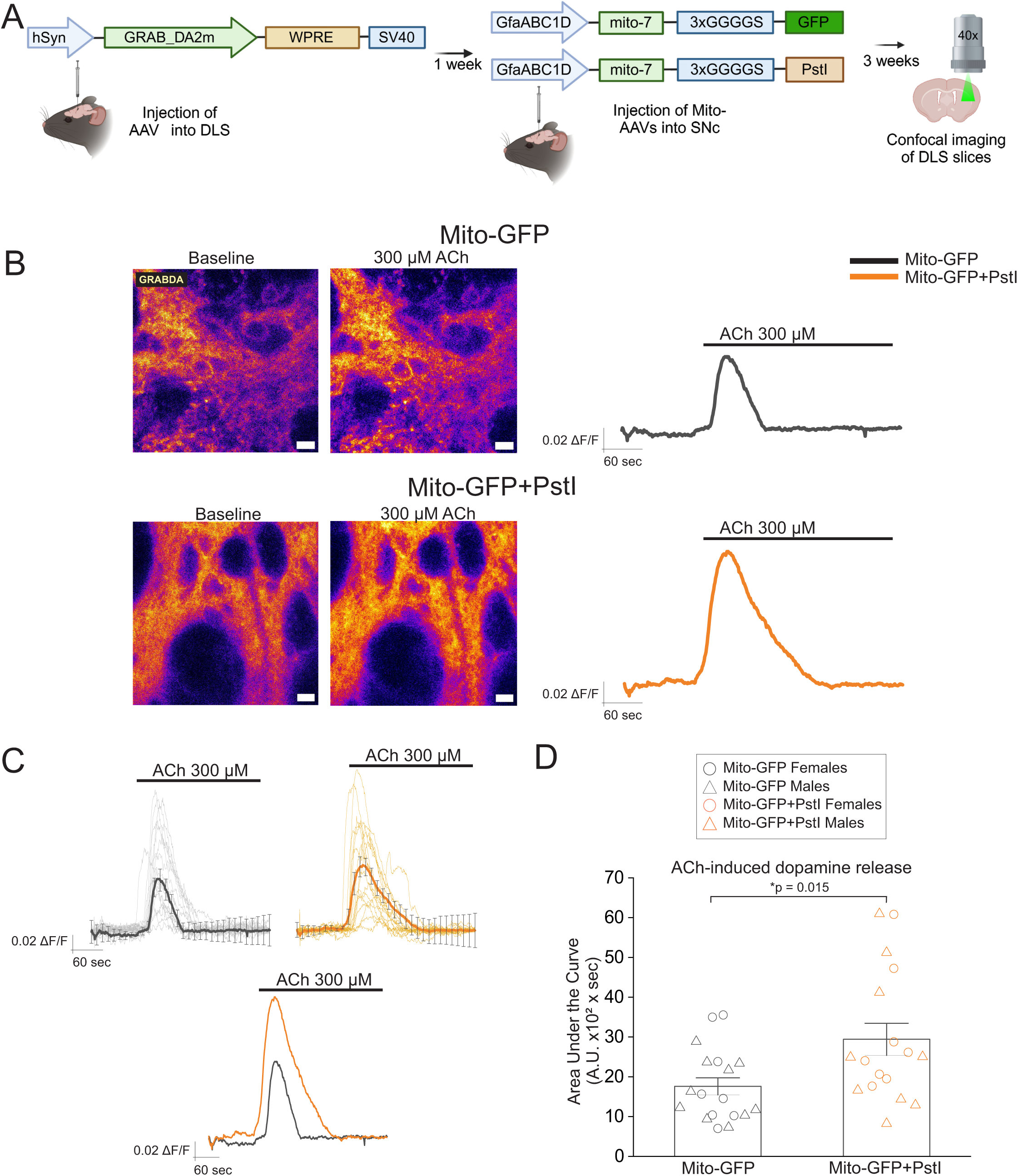
Mito-PstI expression in SNc astrocytes increases ACh-evoked dopamine release by SNc DA neuron terminals within the DLS. (**A**) Schematic showing the experimental timeline. An AAV expressing the optogenetic sensor for dopamine (AAV9-hsyn-GRAB_DA2m) was stereotaxically injected into the DLS of mice one week prior to SNc injection of AAVs expressing either control Mito-GFP or experimental Mito-GFP + Mito-PstI. Acute live brain slices were obtained from these mice and imaged three weeks following SNc AAV injections. Timeline panel was created using BioRender. (**B**) Representative t-stacks (time stacks) of confocal images with pseudo colored Fire look up table (LUT) of GRABDA fluorescence in live DLS brain slices from SNc Mito-GFP (top) and SNc Mito-GFP+PstI (bottom) expressing mice. In each mouse group, left panels show images before ACh bath application and right panels show images during ACh bath application. Scale bar = 10 μm. Traces to the right of each of the panels show average GRABDA fluorescence intensity during bath application of ACh for Mito-GFP (dark grey) and Mito-GFP + Mito-PstI-injected mice (orange) respectively. (**C**) Individual traces per FOV, along with average traces across all FOVs are shown for changes in GRABDA fluorescence intensity during bath application of ACh in Mito-GFP expressing mice (traces per FOV are shown in light grey and average is shown in dark grey) and Mito-GFP + Mito-PstI expressing mice (traces per FOV are shown in light orange and average is shown in dark orange) brain slices. For direct comparison, superimposed average traces of control Mito-GFP in grey and experimental Mito-GFP + Mito-PstI injected mice in orange are also shown. (**D**) Bar graph showing the average AUC with bath application of ACh. Each data point represents a single FOV. Data expressed as mean ± SEM, p-values are based on a Mann Whitney test. n = 18 FOVs from 7 mice for Mito-GFP and n = 17 FOVs from 6 mice for Mito-GFP + Mito-PstI. Individual FOV data points from male and female mice for each condition are shown as circles for females and triangles for males.

### Mito-PstI expression in SNc astrocytes increases spontaneous and apomorphine-induced contralateral rotations in mice

Our observation that Mito-PstI expression in SNc astrocytes dramatically increases ACh-evoked dopamine release within the DLS (Fig. 3) suggests the possibility that mice with Mito-PstI expression in the SNc might also possess abnormally increased dopamine release in the DLS *in vivo*. Based on this rationale, we asked if mtDNA damage due to Mito-PstI expression in SNc astrocytes causes rotational behavior in mice due to imbalances in either dopamine release or D1 and D2 dopamine receptor activation within the DLS. The SNc of mice was injected with AAVs expressing either control Mito-GFP or experimental Mito-GFP + Mito-PstI and then assessed for spontaneous as well as apomorphine-induced rotations in a direction contralateral to the side of AAV injections. Spontaneous and apomorphine-induced rotational assays detect imbalances in dopamine release and / or changes in dopamine receptor sensitivity in the striatum, leading to abnormal rotational behavior in mice in a direction that is contralateral to the side with either excessive striatal dopamine release and / or abnormally increased postsynaptic D1 and D2 dopamine receptor activation (Zarate et al., 2025). An increase in contralateral rotations could be caused by either abnormal presynaptic axonal dopamine release by SNc DA neurons (Konieczny, Lenda, & Czarnecka, 2016) and / or hypersensitive postsynaptic D1 / D2 dopamine receptors (Creese, Burt, & Snyder, 1977; Groppetti et al., 1986; Rouillard, Bedard, Falardeau, & Dipaolo, 1987). Following baseline, day −7 assessment of spontaneous and apomorphine (D1 and D2 dopamine receptor agonist)-induced contralateral rotational behavior prior to SNc AAV injections, on day 0, the SNc of mice were unilaterally injected with AAVs expressing either Mito-GFP (control mice) or Mito-GFP + Mito-PstI (experimental mice), and contralateral rotational behavior was assessed on days 21, 28, 35 and 42 after unilateral AAV injections into the SNc (Fig. 4A).

**Figure 4.**
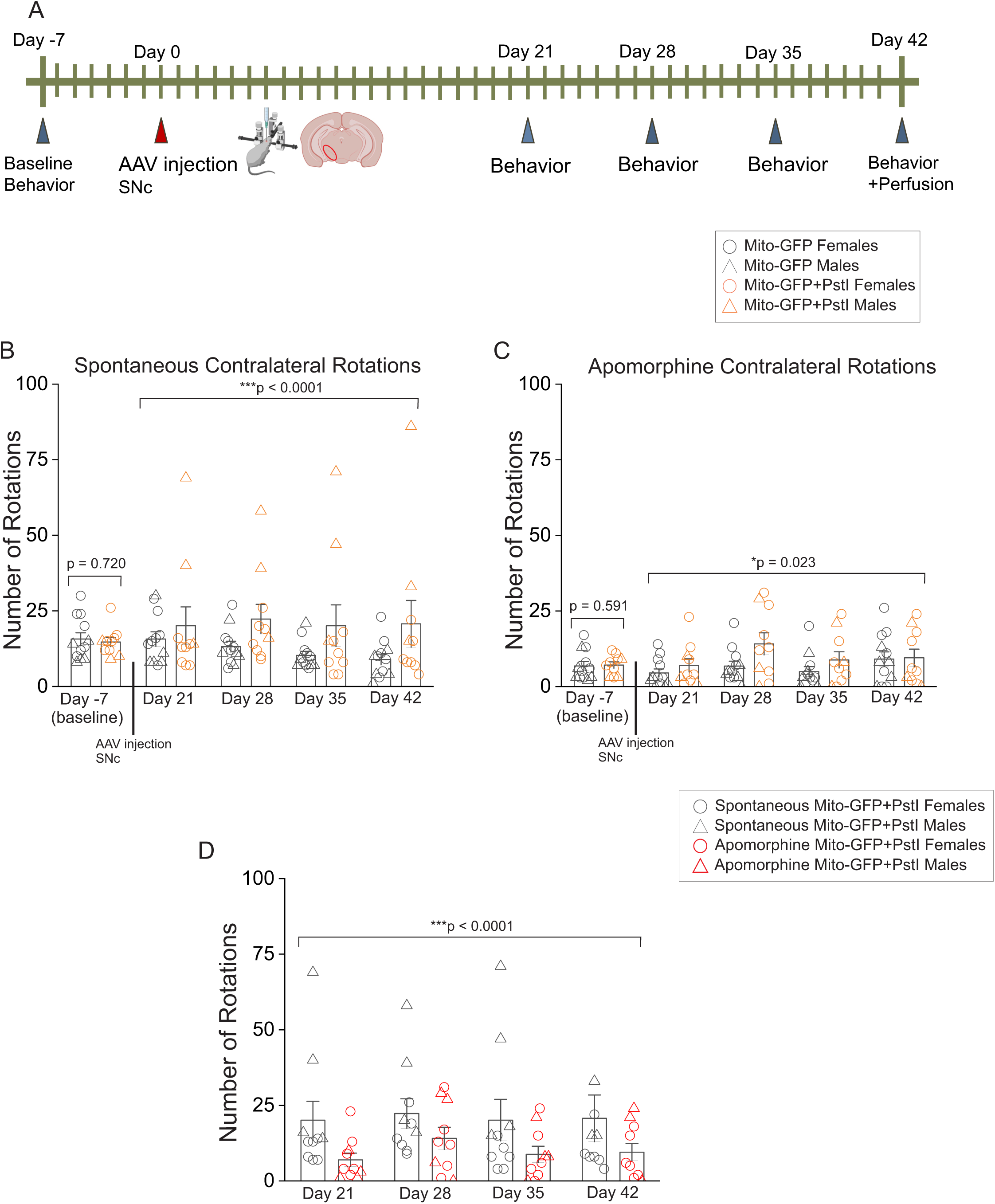
Mito-PstI expression in the SNc increases spontaneous contralateral rotations in mice. (**A**) A schematic of the experimental protocol is shown. Spontaneous and apomorphine-induced contralateral rotational behavior was assessed on days −7 (baseline, prior to AAV injections), and on days 21, 28, 35 and 42 after AAV injections of either Mito-GFP (control group) or Mito-GFP + Mito-PstI (experimental group) into the SNc on day 0. All mice were perfused and their brains extracted immediately after behavioral assessment on day 42 (**B**) Bar graph showing spontaneous contralateral rotations on all days in male and female control Mito-GFP and experimental Mito-GFP + Mito-PstI SNc-injected mice (**C**) Bar graph showing apomorphine-induced contralateral rotations on all days in male and female control Mito-GFP and experimental Mito-GFP + Mito-PstI SNc-injected mice (**D**) Bar graph comparing spontaneous versus apomorphine-induced contralateral rotations on days 21, 35 and 42, after SNc-injection of experimental Mito-GFP + Mito-PstI AAVs into mice. Each data point represents a single mouse. Data expressed as mean ± SEM, *p*-values show main effect of treatment with AAVs or apomorphine based on three way ANOVA and *p*-values comparing two conditions in (**B**) and (**C**) are based on two-sample t-tests. n = 12 mice for Mito-GFP and n = 10 mice for Mito-GFP + Mito-PstI. For all graphs, rotational data in individual mice for each condition are shown as either circles for females or triangles for males.

We found no difference in baseline spontaneous rotations on day −7 between assigned Mito-GFP and Mito-GFP + Mito-PstI mouse groups (Fig. 4B) (Number of contralateral rotations in 10 min: day −7, Mito-GFP = 15.67 ± 2.07 and Mito-GFP + Mito-PstI = 14.7 ± 1.51, *p* = 0.72; two sample t-test). On days 21, 28, 35 and 42, contralateral spontaneous rotations in experimental mice co-expressing Mito-GFP + Mito-PstI in the SNc were significantly increased when compared to control mice expressing Mito-GFP in the SNc (Fig. 4B) (Number of contralateral rotations in 10 min: day 21, Mito-GFP = 15.67 ± 2.38 and Mito-GFP + Mito-PstI = 20.1 ± 6.21; day 28, Mito-GFP = 13.17 ± 1.78 and Mito-GFP + Mito-PstI = 20.1 ± 6.21; day 35, Mito-GFP = 10.25 ± 1.37 and Mito-GFP + Mito-PstI = 22.3 ± 4.86; day 42, Mito-GFP = 8.92 ± 1.77 and Mito-GFP + Mito-PstI = 20.7 ± 7.75; F(1, 72) = 18.59, *p* < 0.0001 with main effect of AAV treatment, 3-way ANOVA).

Apomorphine administration on day −7 did not cause any baseline differences in contralateral rotations between Mito-GFP and Mito-GFP + PstI assigned groups prior to AAV injections (Fig. 4C) (Number of contralateral rotations in 10 min: day −7, Mito-GFP assigned group = 6.83 ± 1.42 and Mito-GFP + Mito-PstI assigned group = 7.1 ± 1.05, *p* = 0.591, two sample t-test). However, the D1 / D2 receptor agonist, apomorphine caused a significant increase in contralateral rotations on days 21, 28, 35 and 42 after AAV injections in the Mito-GFP + Mito-PstI injected mice when compared to control Mito-GFP injected mice (Fig. 4C) (Number of contralateral rotations in 10 min: day 21, Mito-GFP = 4.5 ± 1.34 and Mito-GFP + Mito-PstI = 7 ± 2.20; day 28, Mito-GFP = 6.75 ± 1.64 and Mito-GFP + Mito-PstI = 14.1 ± 3.66; day 35, Mito-GFP = 4.92 ± 1.64 and Mito-GFP + Mito-PstI = 8.8 ± 2.69; day 42, Mito-GFP = 9.08 ± 2.39 and Mito-GFP + Mito-PstI = 9.5 ± 2.86; F(1, 72) = 5.40, p = 0.023, *p* < 0.023, main effect of treatment with AAVs, 3-way ANOVA).

As a further level of analysis, we compared spontaneous versus apomorphine-induced contralateral rotations in experimental mice expressing Mito-GFP + Mito-PstI in the SNc. We found that on all days post-AAV Mito-GFP + Mito-PstI injection into the SNc (days 21, 28, 35 and 42), contralateral rotations were significantly reduced with apomorphine when compared to spontaneous rotations in the same mice (Fig 4D) (Number of contralateral rotations in 10 min for Mito-GFP +Mito-PstI mice: day 21, spontaneous = 20.1 ± 6.21 and apomorphine = 7 ± 2.20; day 28, spontaneous = 22.3 ± 4.86 and apomorphine = 14.1 ± 3.66; day 35, spontaneous = 20.1 ± 6.88 and apomorphine = 8.8 ± 2.69; day 42, spontaneous = 20.7 ± 7.75 and apomorphine = 9.5 ± 2.86; F(1, 64) = 18.31, *p* < 0.0001, main effect of treatment with apomorphine, 3-way ANOVA). Together, these data show that Mito-PstI expression in SNc astrocytes causes an abnormal increase in spontaneous as well as apomorphine-induced contralateral rotational behavior in mice. Furthermore, our data in Figure 4D that directly compare spontaneous versus apomorphine-induced contralateral rotations in Mito-PstI expressing mice suggest that rather than a postsynaptic induction of hypersensitive D1 / D2 dopamine receptors in DLS medium spiny neurons (MSNs) due to Mito-PstI expression in SNc astrocytes, it is likely that abnormal *in vivo* presynaptic increases in dopamine release within the DLS on the side of the SNc injected with Mito-PstI underlie the observed increase in contralateral rotations in these mice.

### Mito-PstI expression in SNc astrocytes is not sufficient to cause a loss of SNc DA neurons

Our data thus far show that Mito-GFP + Mito-PstI expression in mouse SNc astrocytes alters the morphology of astrocytic mitochondria (Fig. 1), causes an abnormal increase in ACh-evoked dopamine release in the DLS (Fig. 3), and significantly increases spontaneous as well as apomorphine-induced contralateral rotations in mice (Fig. 4). Based on these findings, we sought to assess if Mito-PstI expression in SNc astrocytes alone is sufficient to cause a loss of SNc DA neurons. A randomly selected subset of mice subjected to rotational assays in Figure 4 were perfused, and midbrain coronal sections from these mice were stained for TH to visualize SNc DA neurons within the AAV-injected and uninjected SNc of mice (Fig. 5A and B). We employed three independent quantitative readouts of SNc DA neuron loss in TH labeled midbrain sections: (**i**) The number of TH+ SNc DA neurons, (**ii**) TH labeled area in the SNc, and (**iii**) The intensity of SNc TH fluorescence. For all sections from each mouse, the three readouts were applied to both injected and uninjected SNc of control Mito-GFP and experimental Mito-GFP + Mito-PstI expressing mice. This approach enabled us to extract injected / uninjected SNc ratios for each of the three readouts as an additional method to control for differences in antibody staining as well as confocal imaging-related artifacts across multiple midbrain sections.

**Figure 5.**
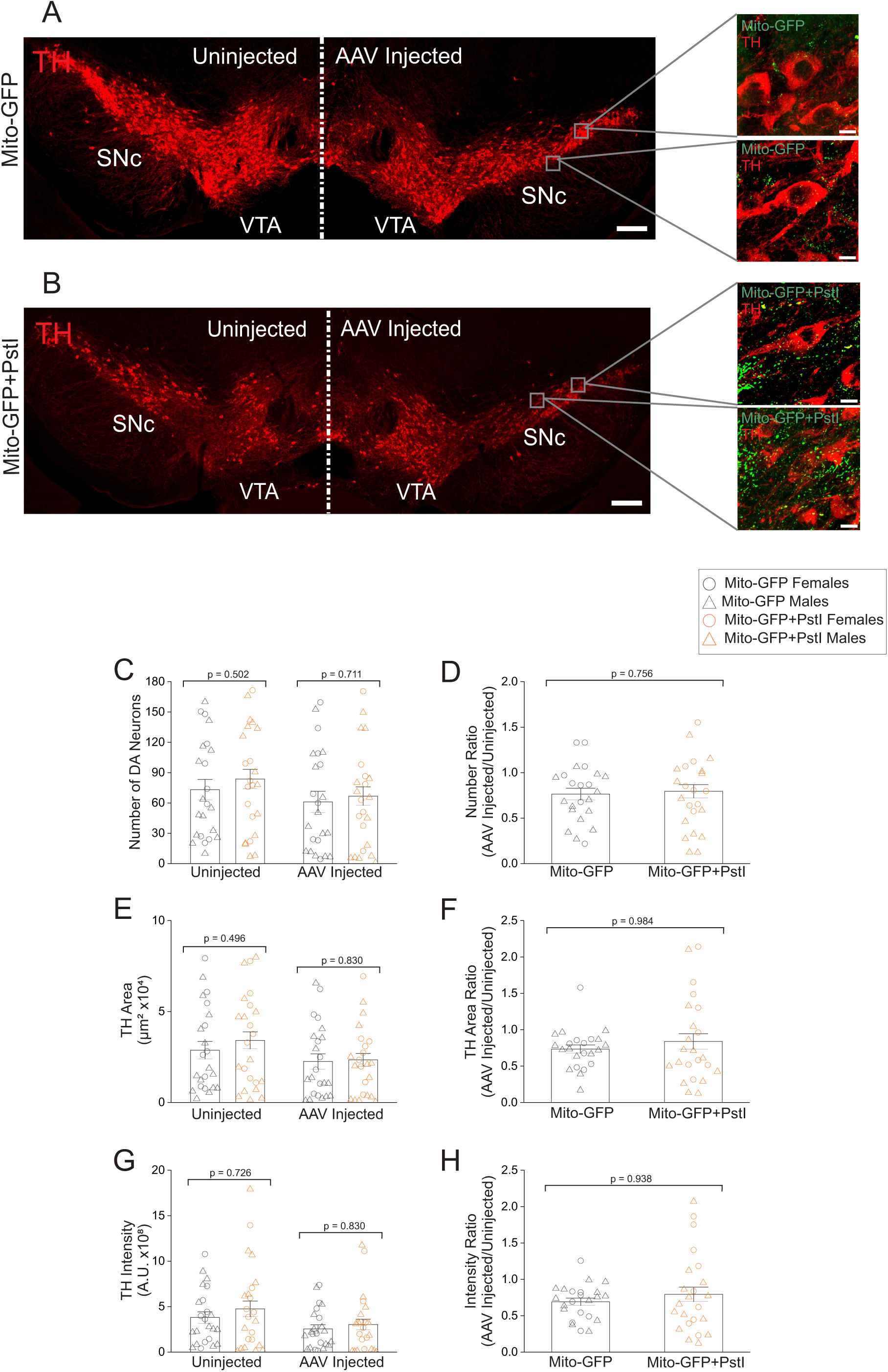
Mito-PstI expression in SNc astrocytes does not cause a loss of SNc DA neurons. (**A**) Representative slide scanner image of a TH labeled midbrain section with unilateral injection of control Mito-GFP AAV into the SNc. Insets on the injected side of the SNc are confocal zoomed in FOVs from the AAV-injected side of the SNc showing GFP expression in SNc astrocytic mitochondria surrounding TH+ SNc DA neurons (**B**) Representative slide scanner image of a TH labeled midbrain section with unilateral injection of experimental Mito-GFP + Mit-PstI AAVs into the SNc. Insets on the injected side of the SNc are confocal zoomed in FOVs from the AAV-injected side of the SNc showing GFP expression in SNc astrocytic mitochondria surrounding TH+ SNc DA neurons. Scale bar = 200 μm for slide scanner image of midbrain sections and 10 μm for zoomed in confocal insets (**C**) Bar graph showing number of DA neurons in the SNc for AAV-injected and uninjected sides of the SNc in control Mito-GFP and experimental Mito-GFP + Mito-PstI SNc-injected mice (**D**) Bar graph showing ratios of number of DA neurons between the SNc AAV-injected and uninjected sides of the SNc for control Mito-GFP and experimental Mito-GFP + Mito-PstI SNc-injected mice (**E**) Bar graph showing area of TH+ immunolabeling in the SNc for AAV-injected and uninjected sides of the SNc in control Mito-GFP and experimental Mito-GFP + Mito-PstI SNc-injected mice (**F**) Bar graph showing ratios of area of TH+ immunolabeling between the SNc AAV-injected and uninjected sides of the SNc for control Mito-GFP and experimental Mito-GFP + Mito-PstI SNc-injected mice (**G**) Bar graph showing integrated intensity of TH+ immunolabeling in the SNc for AAV-injected and uninjected sides of the SNc in control Mito-GFP and experimental Mito-GFP + Mito-PstI SNc-injected mice (**H**) Bar graph showing ratios of integrated intensity of TH+ immunolabeling between the SNc AAV-injected and uninjected sides of the SNc for control Mito-GFP and experimental Mito-GFP + Mito-PstI SNc-injected mice. Each data point represents one midbrain section. Data expressed as mean ± SEM, *p*-values are based on a two-sample t-test. n = 23 sections from 7 mice for Mito-GFP and n = 27 sections from 8 mice for Mito-GFP + Mito-PstI. Individual data points for male and female mice are shown as circles for females and triangles for males.

There was no significant difference between control Mito-GFP and experimental Mito-GFP + PstI mice in the number of TH+ SNc DA neurons in both AAV-injected and uninjected sides (Fig. 5C) (Number of TH+ SNc DA neurons: Uninjected Mito-GFP = 71.77 ± 15.61 and Uninjected Mito-GFP + Mito-PstI = 83.47 ± 15.97, *p* = 0.611, two-sample t-test; Injected Mito-GFP = 57.88 ± 16.63 and Injected Mito-GFP + Mito-PstI = 66.99 ± 14.81, *p* = 0.688, two-sample t-test). There was also no significant difference in TH labeled neuron area between Mito-GFP and Mito-GFP + PstI mice (Fig. 5E) (Area of TH+ SNc DA neurons in μm^2^: Uninjected Mito-GFP = 25696.77 ± 6498.95 and Uninjected Mito-GFP + Mito-PstI = 32798.22 ± 7952.24, *p* = 0.502, two-sample t-test; Injected Mito-GFP = 21788.61 ± 6828.54 and Injected Mito-GFP + Mito-PstI = 23531.17 ± 5380.43, *p* = 0.842, two-sample t-test). Finally, the intensity of TH+ SNc DA neurons did not show any difference between control Mito-GFP and experimental Mito-GFP + PstI mice in the AAV-injected and uninjected SNc (Fig. 5G) (Intensity of TH+ SNc DA neurons expressed as A.U.: Uninjected Mito-GFP = 3.67×10^8^ ± 1.06×10^8^ and Uninjected Mito-GFP + Mito-PstI = 3.77×10^8^ ± 1×10^8^, *p* = 0.950, two-sample t-test; Injected Mito-GFP = 2.55×10^8^ ± 7.82×10^7^ and Injected Mito-GFP + Mito-PstI = 3.11×10^8^ ± 8.86×10^7^, *p* = 0.646, two-sample t-test).

As a further level of analysis, we obtained ratios between the injected and uninjected SNc for each midbrain section and compared these ratios between control Mito-GFP and experimental Mito-GFP + Mito-PstI mice for each of the three parameters, *viz.,* SNc DA cell number, TH+ area and TH intensity. There were no differences in AAV-Injected / Uninjected SNc ratios between control Mito-GFP and experimental Mito-GFP + Mito-PstI mice for either cell number (Fig. 5D) (Injected / Uninjected side ratio for number of TH+ SNc DA neurons: Mito-GFP = 0.724 ± 0.081 and Mito-GFP + Mito-PstI = 0.786 ± 0.100, *p* = 0.649, two-sample t-test) or TH+ SNc DA neuron area (Fig. 5F) (Injected / Uninjected side ratio for area of TH+ SNc DA neurons: Mito-GFP = 0.734 ± 0.057, Mito-GFP + Mito-PstI: 0.840 ± 0.105, *p* = 0.984, Mann-Whitney test) or SNc DA neuron TH intensity (Fig. 5G) (Injected / Uninjected side ratio for intensity of TH in SNc DA neurons: Mito-GFP = 0.663 ± 0.062 A.U., Mito-GFP + Mito-PstI = 0.683 ± 0.095, *p* = 0.862, two-sample t-test). Taken together, these data show that astrocytic expression of Mito-PstI alone in the SNc was not sufficient to cause a loss of SNc DA neurons.

### Mito-PstI expression in SNc astrocytes increases spontaneous but not apomorphine-induced contralateral rotations in a low dose 6-OHDA mouse model of parkinsonism

Although we did not observe a loss of SNc DA neurons with Mito-PstI expression in SNc astrocytes, our data show that Mito-PstI expression in mouse SNc astrocytes causes SNc DA neurons to become dysfunctional, which is evidenced by abnormal increases in ACh-evoked dopamine release within the DLS (Fig. 3), as well as an increase in spontaneous and apomorphine-induced contralateral rotations in Mito-PstI expressing mice (Fig. 4). Since SNc DA neuron dysfunction is a known predisposing factor for neurodegeneration in PD (Watanabe, Dijkstra, & Nagatsu, 2024), we asked if Mito-PstI expression in SNc astrocytes can worsen parkinsonian motor deficits in mice following injection of a low 3 μg dose of the neurotoxin, 6-OHDA into the mouse DLS. To test this hypothesis, the SNc of mice was injected with either Mito-GFP (control) or Mito-GFP + Mito-PstI (experimental), and three weeks after AAV injections into the SNc, all mice were unilaterally injected with low dose 6-OHDA (3 μg in 1 μl) into the DLS on the same side as SNc AAV injections (Fig. 6A). 6-OHDA is a neurotoxin that induces loss of SNc DA neurons when injected into the DLS by causing a dying back of SNc DA neuron axonal terminals in the DLS (Fig. 6B) (Ichitani, Okamura, Matsumoto, Nagatsu, & Ibata, 1991).

**Figure 6.**
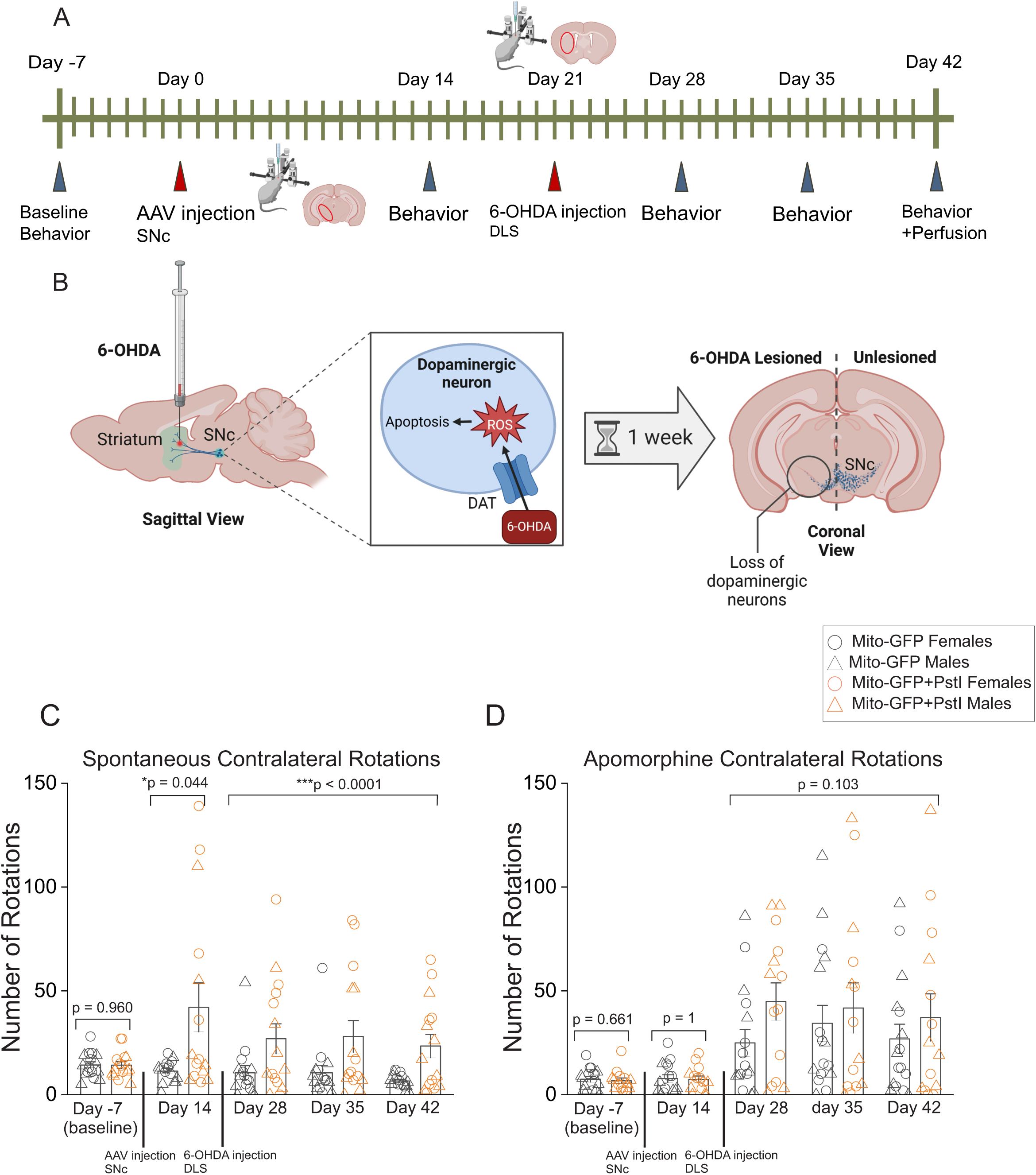
Mito-PstI expression in SNc astrocytes increases spontaneous contralateral rotations in a low dose 6-OHDA mouse model. (**A**) A schematic of the experimental protocol is shown. Spontaneous and apomorphine-induced contralateral rotational behavior was assessed on days −7 (baseline, prior to AAV injections), and on days 28, 35 and 42 after AAV injections of either Mito-GFP (control group) or Mito-GFP + Mito-PstI (experimental group) into the SNc on day 0 and injection of low dose (3 μg) 6-OHDA into the DLS of all mice on day 21. All mice were perfused and their brains extracted immediately after behavioral assessment on day 42 (**B**) A schematic of the model for 6-OHDA-induced SNc DA neuron loss is shown. 6-OHDA was stereotaxically injected into the DLS, where it enters SNc DA neuron axonal terminals through the dopamine transporter (DAT) and causes death of SNc DA neurons via reactive oxygen species (ROS)-induced apoptosis, beginning ~1 week after 6-OHDA injection into the DLS. Panel created in BioRender (**C**) Bar graph showing spontaneous contralateral rotations prior to and following SNc injections of AAV-Mito-GFP or AAV Mito-GFP + Mito-PstI and low dose 6-OHHDA into the DLS. (**D**) Bar graph showing apomorphine-induced contralateral rotations prior to and following SNc injections of AAV-Mito-GFP or AAV Mito-GFP + Mito-PstI and low dose 6-OHHDA into the DLS. Each data point represents a single mouse. Data expressed as mean ± SEM, *p*-values show main effect of treatment with AAVs or apomorphine based on three way ANOVA and *p*-values comparing two conditions in (**C**) and (**D**) are based on two-sample t-tests. n = 16 mice for Mito-GFP and n = 15 mice for Mito-GFP + Mito-PstI. For all graphs, rotational data in individual mice for each condition are shown as either circles for females or triangles for males.

Spontaneous and apomorphine-induced contralateral rotations in mice were longitudinally assessed as follows: (**i**) On day −7 as a baseline prior to any injections, (**ii**) On day 14, after injecting only control Mito-GFP and experimental Mito-GFP + Mito-PstI AAVs into the mouse SNc, and (**iii**) On days 28, 35 and 42, after additionally injecting low dose 6-OHDA into the DLS of the same mice (Fig. 6A and B). We found no difference in baseline day −7 spontaneous rotations between assigned Mito-GFP and Mito-GFP + Mito-PstI mouse groups prior to any treatment (Fig. 6C) (Number of spontaneous contralateral rotations in 10 min: day −7, Mito-GFP = 14.31 ± 1.50 and Mito-GFP + Mito-PstI = 14.2 ± 1.66, *p* = 0.96; two sample t-test). Two weeks after unilateral Mito-GFP or Mito-GFP + Mito-PstI injection into the SNc, we found a significant increase in spontaneous contralateral rotations in experimental Mito-GFP + Mito-PstI mice when compared control Mito-GFP mice (Fig. 6C) (Number of spontaneous contralateral rotations in 10 min: day 14, Mito-GFP = 11 ± 1.39 and Mito-GFP + Mito-PstI = 42.07 ± 11.78, *p* = 0.042; Mann-Whitney test). Following injection of low dose 6-OHDA into the DLS, spontaneous rotations were significantly increased on days 28, 35 and 42 in experimental 6-OHDA + Mito-GFP + Mito-PstI injected mice when compared to control 6-OHDA + Mito-GFP injected mice (Fig. 6C) (Number of contralateral rotations in 10 min in 6-OHDA-injected mice: day 28, Mito-GFP = 10.75 ± 3.24 and Mito-GFP + Mito-PstI = 26.93 ± 7.18; day 35, Mito-GFP = 10.56 ± 3.60 and Mito-GFP + Mito-PstI = 28.07 ± 7.61; day 42, Mito-GFP = 6.56 ± 0.83 and Mito-GFP + Mito-PstI = 23.47 ± 5.62; F(1, 81)= 15.65, *p* < 0.0001 with main effect of AAV treatment, 3-way ANOVA).

Apomorphine-induced contralateral rotations between control Mito-GFP and experimental Mito-GFP + Mito-PstI injected mice showed no baseline differences on day −7 (Fig. 6D) (Number of apomorphine-induced contralateral rotations in 10 min without AAVs and 6-OHDA: day −7, Mito-GFP = 7.44 ± 1.34 and Mito-GFP + Mito-PstI = 6.71 ± 1.33, *p* = 0.661, Mann-Whitney test). Injection of low dose 6-OHDA into DLS significantly increased apomorphine-induced contralateral rotations to an equal extent in control Mito-GFP as well as experimental Mito-GFP + Mito-PstI-injected mice when compared to mice expressing only Mito-GFP or Mito-GFP + Mito-PstI without 6-OHDA (Fig. 6D), suggesting that low dose injection of 6-OHDA into the DLS induced a parkinsonian deficit in both groups of mice unilaterally expressing either control Mito-GFP or experimental Mito-GFP + Mito PstI in the SNc. However, there were no significant differences in apomorphine-induced contralateral rotations between control Mito-GFP and experimental Mito-GFP + Mito-PstI-injected mice on days 28, 35 and 42 after low dose 6-OHDA injection into the ipsilateral DLS (Fig. 6D) (Number of apomorphine-induced contralateral rotations in 10 min in 6-OHDA-injected Mito-GFP mice: day 28 = 24.94 ± 6.46, day 35 = 34.38 ± 8.63, day 42 = 26.88 ± 7.06 and Mito-GFP + Mito-PstI mice: day 28 = 44.86 ± 8.96, day 35 = 41.79 ± 12.09, day 42 = 37.21 ± 11.36; F(1, 78)= 2.73, *p* = 0.103 with main effect of treatment, 3-way ANOVA). Taken together, these data show that astrocytic mtDNA damage induced by Mito-PstI expression in SNc astrocytes in a mouse model of parkinsonism with low dose 6-OHDA injection into the DLS worsens parkinsonian deficits. This is demonstrated by a specific increase in spontaneous contralateral rotations, likely caused by increases in presynaptic dopamine release (Fig. 3). Furthermore, our data showing that apomorphine reduces contralateral rotations in Mito-PstI expressing mice suggests that apomorphine likely overrides Mito-PstI-induced increases in presynaptic dopamine release by directly activating 6-OHDA-induced hypersensitive postsynaptic D1 and D2 receptors in MSNs within the DLS of these mice.

### Mito-PstI expression in the SNc astrocytes exacerbates SNc DA neuron loss in 6-OHDA-treated mice

Since Mito-PstI expression in SNc astrocytes increases spontaneous contralateral rotations in 6-OHDA-treated mice (Fig. 6C), we sought to determine if Mito-PstI worsens the loss of SNc DA neurons following low dose 6-OHDA injection into the DLS. To do this, a randomly selected subset of mice subjected to rotational assays in Figure 6 were perfused, and midbrain coronal sections from these mice were stained for TH to quantify DA neurons within the SNc, expressing either control Mito-GFP or experimental Mito-GFP + Mito-PstI (Fig. 7A and B), with ipsilateral injection of low dose 6-OHDA into the DLS. As described in Figure 5, we used three independent readouts, *viz.*, SNc DA neuron number, TH area and TH intensity to quantify SNc DA neuron loss in these mice. For each of the three readouts, ratios of the 6-OHDA lesioned / unlesioned sides of the SNc were used to compare SNc DA neuron loss between control Mito-GFP and experimental Mito-GFP + Mito PstI expressing mice.

**Figure 7.**
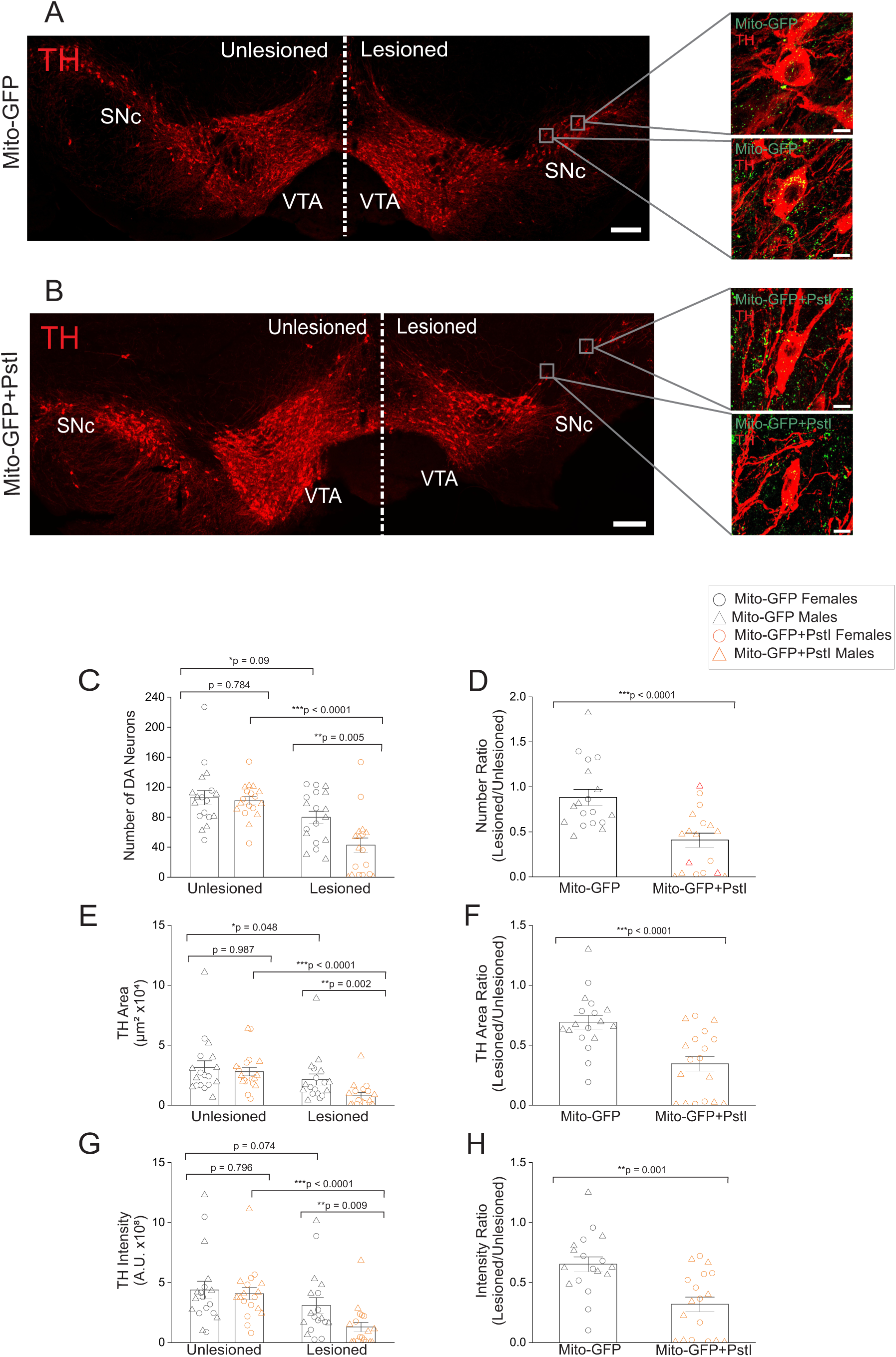
Mito-PstI expression in SNc astrocytes exacerbates the loss of SNc DA neurons following low dose injection of 6-OHDA into the DLS. (**A**) Representative slide scanner image of a TH labeled midbrain section with unilateral injection of control Mito-GFP AAV into the SNc and ipsilateral 6-OHDA injection into the DLS. Insets on the injected side of the SNc are confocal zoomed in FOVs from the SNc AAV + DLS 6-OHDA-injected side showing GFP expression in SNc astrocytic mitochondria surrounding TH+ SNc DA neurons (**B**) Representative slide scanner image of a TH labeled midbrain section with unilateral injection of experimental Mito-GFP + Mit-PstI AAVs into the SNc and ipsilateral 6-OHDA injection into the DLS. Insets on the injected side of the SNc are confocal zoomed in FOVs from the AAV + DLS 6-OHDA-injected side showing GFP expression in SNc astrocytic mitochondria surrounding TH+ SNc DA neurons. Scale bar = 200 μm for slide scanner image of midbrain sections and 10 μm for zoomed in confocal insets (**C**) Bar graph showing number of DA neurons in the SNc of SNc AAV + DLS 6-OHDA-injected and uninjected sides for control Mito-GFP and experimental Mito-GFP + Mito-PstI SNc-injected mice (**D**) Bar graph showing ratios of number of DA neurons in the SNc between the SNc AAV + DLS 6-OHDA-injected and uninjected sides for control Mito-GFP and experimental Mito-GFP + Mito-PstI SNc-injected mice (**E**) Bar graph showing area of TH+ immunolabeling in the SNc for SNc AAV + DLS 6-OHDA-injected and uninjected sides of the SNc for control Mito-GFP and experimental Mito-GFP + Mito-PstI SNc-injected mice (**F**) Bar graph showing ratios of area of TH+ immunolabeling between the SNc AAV + DLS 6-OHDA-injected and uninjected sides of the SNc for control Mito-GFP and experimental Mito-GFP + Mito-PstI SNc-injected mice (**G**) Bar graph showing integrated intensity of TH+ immunolabeling in the SNc for SNc AAV + DLS 6-OHDA-injected and uninjected sides for control Mito-GFP and experimental Mito-GFP + Mito-PstI SNc-injected mice (**H**) Bar graph showing ratios of integrated intensity of TH+ immunolabeling in the SNc between the SNc AAV + DLS 6-OHDA-injected and uninjected sides for control Mito-GFP and experimental Mito-GFP + Mito-PstI SNc-injected mice. Each data point represents one midbrain section. Data expressed as mean ± SEM, *p*-values are based on a two sample t-test. n = 18 sections from 6 mice for Mito-GFP and n = 19 sections from 6 mice for Mito-GFP + Mito-PstI. Individual data points for male and female mice are shown as circles for females and triangles for males.

The number of SNc DA neurons in the uninjected side was similar between control Mito-GFP and experimental Mito-GFP + Mito-PstI mouse groups (Fig. 7C). For both groups of mice, when compared to the 6-OHDA unlesioned side, low dose 6-OHDA injection into the DLS caused a significant decrease in SNc DA neuron number only on the side injected with 6-OHDA + SNc AAVs (Mito-GFP or Mito-GFP + Mito-PstI), indicating successful 6-OHDA-induced retrograde loss of SNc DA neurons (Fig. 6B and 7C). Importantly, we observed a dramatic ~50% greater decrease in the number of SNc DA neurons in experimental mice expressing Mito-GFP + Mito-PstI in SNc astrocytes in the DLS 6-OHDA lesioned side when compared to the DLS 6-OHDA lesioned SNc of control Mito-GFP expressing mice (Fig. 7C) (Number of TH+ SNc DA neurons per section: Unlesioned, no Mito-GFP = 106.056 ± 9.522 and Unlesioned, no Mito-GFP + Mito-PstI = 102.026 ± 5.293, *p* = 0.784, Mann-Whitney test; 6-OHDA lesioned + Mito-GFP = 79.889 ± 7.956 and 6-OHDA lesioned + Mito-GFP + Mito-PstI = 42.684 ± 9.554, *p* = 0.005, Mann-Whitney test).

We also observed a 6-OHDA-induced decrease in SNc TH area as well as TH intensity on the 6-OHDA lesioned side when compared to the unlesioned side for both control Mito-GFP and experimental Mito-GFP + Mito-PstI SNc injected mice. Similar to our finding with TH+ SNc DA neuron numbers, there was a dramatic ~60 % decrease in TH area as well as TH intensity in experimental Mito-GFP + Mito-PstI SNc injected mice in the 6-OHDA lesioned side when compared to control Mito-GFP SNc injected mice in the 6-OHDA lesioned side (Fig 7E and G) (Area of TH+ SNc DA neurons: Unlesioned, no Mito-GFP = 31395.52 ± 5606.31 μm^2^ and Unlesioned, no Mito-GFP + Mito-PstI = 28000.19 ± 3506.35 μm^2^, *p* = 0.987, Mann-Whitney test; 6-OHDA lesioned + Mito-GFP = 21441.29 ± 4572.84 μm^2^ and 6-OHDA lesioned + Mito-GFP + Mito-PstI = 8273.52 ± 2222.83 μm^2^, *p* = 0.002, Mann-Whitney test; Intensity of TH+ SNc DA neurons in A.U: Unlesioned, no Mito-GFP = 4.38×10^8^ ± 7.26×10^7^ and Unlesioned, no Mito-GFP + Mito-PstI = 4.08×10^8^ ± 4.95×10^7^, *p* = 0.796, Mann-Whitney test; 6-OHDA lesioned + Mito-GFP = 3.10×10^8^ ± 6.50×10^7^ and 6-OHDA lesioned + Mito-GFP + Mito-PstI = 1.29×10^8^ ± 3.75×10^7^, *p* = 0.009, Mann-Whitney test).

Ratios of the lesioned and unlesioned SNc for TH+ SNc DA neuron number, area and intensity between Mito-GFP and Mito-GFP + Mito-PstI-injected mice showed similar ~50 % decreases in experimental Mito-GFP + Mito-PstI-injected mice when compared to control Mito-GFP-injected mice for each of the 3 readouts (Fig. 7D, F and H) (Ratio of number of TH+ SNc DA neurons: Mito-GFP = 0.882 ± 0.088 and Mito-GFP + Mito-PstI = 0.408 ± 0.078, *p* < 0.0001, Mann-Whitney test; Ratio of area of TH+ SNc DA neurons: Mito-GFP = 0.692 ± 0.058 and Mito-GFP + Mito-PstI = 0.345 ± 0.062, *p* < 0.0001, Mann-Whitney test; Ratio of intensity of TH+ SNc DA neurons: Mito-GFP = 0.653 ± 0.061 and Mito-GFP + Mito-PstI = 0.319 ± 0.060, *p* = 0.001, Mann-Whitney test). Taken together, these data clearly show that Mito-PstI expression in SNc astrocytes exacerbates SNc DA neuron loss in a low dose 6-OHDA mouse model of parkinsonism.

## Discussion

In this study, we use our recently developed AAV-based tool called Mito-PstI (Ayala et al., 2024) to assess the effect of SNc astrocytic mtDNA damage on mouse DA neuron function and viability *in vivo*. Experiments examining the effect of Mito-PstI on SNc astrocytic mitochondria and astrocyte morphology highlight three specific points: (**i**) Control Mito-GFP labeling of astrocytic mitochondria in the SNc reveal isolated mitochondrial particles in close vicinity to TH+ SNc DA neuron soma (Fig. 1A and E). This distinct morphology is in stark contrast to the fusiform morphology of mitochondria previously reported by us in DLS astrocytes (Ayala et al., 2024; Huntington & Srinivasan, 2021). These data further demonstrate that astrocytic mitochondria in central nervous system astrocytes can be morphologically heterogenous, perhaps depending on energy demands of the brain region in which they reside. (**ii**) Mito-PstI expression in the SNc causes a nearly 50 % increase in the number and area of astrocytic mitochondria in the SNc (Fig. 1C and D). We infer that this is the result of mtDNA damage causing alterations in mitochondrial dynamics, which can affect mitophagy and mitochondrial biogenesis (Rong et al., 2021; Zhao & Sumberaz, 2020). Our conclusion on the effects on mitochondrial dynamics is limited by confocal resolution (~0.45 μm), which prevents determination of changes in individual mitochondrial particle size below this limit following Mito-PstI expression in SNc astrocytes. (**iii**) Mito-PstI-induced astrocytic mtDNA damage in the SNc is accompanied by a generalized increase in astrocytic process density and a significant increase in the wrapping of processes around TH+ SNc DA neurons (Fig. 2C and D). Although morphological changes in astrocytes following mtDNA damage have been previously reported in relation to increased astrocyte reactivity (Ignatenko et al., 2018), we observe a paradoxical decrease in S100B intensity (Fig. 2E). This suggests that the observed changes in SNc astrocyte processes are not due to an increase in astrocyte reactivity, but rather, a hitherto unknown effect of SNc astrocytic mtDNA damage on S100B expression. Indeed, studies show that human induced pluripotent stem cell (iPSC)-derived astrocytes with a G2019S PD-associated point mutation in the leucine-rich repeat kinase 2 (LRRK2) protein demonstrate a decrease in S100B expression (Ramos-Gonzalez et al., 2021), and that LRRK2 G2019S mutations in human iPSCs cause mtDNA damage (Sanders et al., 2014). These reports, in conjunction with our data lend credence to the idea that astrocytic mtDNA damage can attenuate S100B expression in astrocytes.

In further experiments, we show that Mito-PstI expression in SNc astrocytes causes a robust increase in ACh-evoked dopamine release from SNc axonal terminals within the DLS (Fig. 3D). One mechanistic explanation for this result is that the increase in wrapping and density of astrocytic processes around the soma of SNc DA neurons by Mito-PstI-expressing astrocytes (Fig. 2) alters autoregulatory pathways governing dopamine release in the DLS. In this regard, previous studies have shown that the release of dopamine by SNc DA neurons can act on somatodendritic D2 receptors, thereby inhibiting dopamine release at DA axonal terminals in the striatum (Cragg & Greenfield, 1997; Hikima et al., 2021; Sulzer, Cragg, & Rice, 2016). Therefore, we surmise that increased wrapping of DA neurons by Mito-PstI expressing astrocytic processes in the SNc could reduce extracellular dopamine levels in the SNc via enhanced astrocytic reuptake of released dopamine in the SNc, thereby altering ACh-evoked dopamine release in DA axonal terminals within the DLS. In support of the idea that physical astrocytic coverage of neural structures can alter neurotransmitter release, multiple studies have shown that changes in astrocyte coverage within neural structures can affect the concentration of extrasynaptic neurotransmitters, thereby modulating the function of nearby neurons (Lawal, Ulloa Severino, & Eroglu, 2022). Based on this rationale, we suggest the possibility that expression of Mito-PstI in the SNc disrupts D2 receptor-mediated SNc somatodendritic DA neuron autoregulation of axonal dopamine due to reduced levels of extracellular dopamine in the SNc. Dysregulated dopamine homeostasis in SNc DA neurons could in turn cause an abnormal increase in the releasable pool of dopamine vesicles at axonal terminals within the DLS via alterations in axonal vesicular dopamine release probability and / or reuptake of released dopamine at axonal terminals of SNc DA neurons within the DLS (Jin et al., 2024; Sulzer et al., 2016). It should be noted that the observed increase in ACh-evoked dopamine release within the DLS likely does not involve an increase in dopamine synthesis because expression of Mito-PstI in SNc astrocytes has no effect on TH expression (Fig. 5C), which is a rate limiting enzyme for dopamine synthesis.

Our novel finding that Mito-PstI expression in the SNc induces an increase in ACh-evoked dopamine release by axonal terminals within the DLS (Fig. 3) is supported by behavioral data showing that when compared to control Mito-GFP-injected mice, unilateral Mito-PstI expression in SNc astrocytes alone is sufficient to cause a significant increase in spontaneous as well as apomorphine-induced contralateral rotations across multiple weeks (Fig. 4B and C). Interpretation of this observation is based on the reasoning that unilateral damage to the SNc can cause contralateral rotations in mice via postsynaptic and / or presynaptic mechanisms. A known postsynaptic mechanism is that after unilateral loss of SNc DA neurons, apomorphine activates hypersensitive postsynaptic D1 and D2 dopamine postsynaptic receptors in MSNs within the DLS on the SNc damaged side, which results in apomorphine-induced contralateral rotations (Zarate et al., 2025). By contrast, a presynaptic mechanism could depend on excessive dopamine release by axonal terminals of SNc neurons in the DLS on the side with SNc damage, which can also similarly result in increased contralateral rotational behavior (Konieczny et al., 2016). To parse out whether pre- or post-synaptic mechanisms underlie the observed increases in spontaneous and apomorphine-induced contralateral rotations with Mito-PstI-induced mtDNA damage in SNc astrocytes, we directly compared spontaneous contralateral rotational data to apomorphine-induced contralateral rotations in mice expressing Mito-PstI in the SNc (Fig. 4D). In this context, our finding that apomorphine reduces contralateral rotations in mice with SNc expression of Mito-PstI (Fig. 4D), when taken in conjunction with our data showing increased ACh-evoked dopamine release in live DLS slices from SNc Mito-PstI expressing mice (Fig. 3), suggests that Mito-PstI expression in the SNc likely causes an abnormal increase in presynaptic dopamine release by SNc axonal terminals in the DLS *in vivo*, which significantly increases contralateral rotations in mice expressing Mito-PstI in SNc astrocytes.

Despite our observation that Mito-PstI expression in SNc astrocytes causes an abnormal increase in ACh-evoked dopamine release in the DLS (Fig 3), as well as an increase in contralateral rotational behavior (Fig. 4), these mice do not show an increase in SNc DA neuron loss when compared to control mice expressing Mito-GFP in the SNc (Fig. 5). Based on these data, we rationalize that mtDNA damage to SNc astrocytes could act as an additional insult during ongoing parkinsonian pathology, thereby worsening SNc DA neuron loss. In agreement with our rationale, we show that in mice with low dose 6-OHDA-induced parkinsonism, Mito-PstI expression in the SNc not only causes a persistent increase in spontaneous contralateral rotations across multiple weeks (Fig. 6C) but also results in an ~50% greater loss of DA neurons in the SNc (Fig. 7). It is notable that apomorphine-induced contralateral rotations in experimental Mito-PstI mice are not significantly different from control Mito-GFP mice (Fig. 6D). This result suggests that the D1 / D2 receptor agonist, apomorphine activates 6-OHDA-induced hypersensitive postsynaptic D1 and D2 receptors in DLS MSN neurons, which could override the rotational behavior caused by increased dopamine release within the DLS of 6-OHDA treated Mito-PstI mice. On the other hand, spontaneous contralateral rotations in 6-OHDA treated Mito-PstI expressing mice remain significantly elevated, likely because of SNc Mito-PstI-induced increases in presynaptic dopamine release that activate hypersensitive MSN D1 and D2 receptors *in vivo*.

For the first time, this study shows that mtDNA damage, specifically in SNc astrocytes causes an abnormal increase in dopamine release within the striatum, leading to an exacerbation of SNc DA neuron loss. Given the multiple lines of evidence presented here, demonstrating an abnormal increase in dopamine release in the DLS following astrocytic mtDNA damage by Mito-PstI in the SNc (Figs. 3, 4, 6 and 7), we infer that mtDNA damage in SNc astrocytes worsens parkinsonian behavioral deficits and SNc DA neuron loss by abnormally increasing presynaptic dopamine release at SNc DA neuron axonal terminals within the DLS. In support of this view, the established Thy1-α-synuclein mouse model of PD demonstrates an abnormal increase in striatal dopamine release prior to SNc DA neurodegeneration at 4 months (Lam et al., 2011), and overexpression of TH in mice, which is the rate limiting enzyme for dopamine synthesis results in hyperoxidative stress, which can eventually lead to accelerated SNc DA neuron loss (Vecchio et al., 2021). Together, these reports suggest that sustained increases in dopamine release within the striatum could result in the toxic accumulation of hyperoxidative dopamine metabolites such as reactive oxygen species and dopamine quinones, causing a dying back of SNc DA axons in the striatum and thereby leading to the accelerated loss of SNc DA neurons.

In summary, this study positions SNc astrocytic mtDNA damage as an important component to consider in our understanding of the role of astrocytes in PD pathogenesis. From a translational perspective, recent studies have shown that patients with clinical PD possess an increased accumulation of mtDNA mutations in blood cells (Qi et al., 2023; Tresse et al., 2023), which invokes the intriguing view that an accumulation of mtDNA mutations in CNS astrocytes could contribute to the progression of clinical PD. In this context, our study demonstrates that mtDNA damage in SNc astrocytes could indeed cause SNc DA neuron dysfunction, thereby worsening neurodegeneration during PD. Taken together, our findings highlight the potential of targeting mtDNA damage in SNc astrocytes as a clinically relevant disease-modifying strategy against PD.

## Acknowledgements

The authors acknowledge the assistance of the Integrated Microscopy and Imaging Laboratory at the Texas A&M College of Medicine. RRID:SCR_021637. This work was funded by a research grant from the National Institutes of Health (NIH)/NINDS, R21 NS121959 to MR and RS. DA was partially funded by a NIH T32 GM135748 grant.

## Author contributions

DAA performed all experiments, analyzed data and created the figures. GMH analyzed data related to mitochondrial morphology and astrocyte reactivity. AA, ZK, LL, DP and ET performed experiments and analyzed data. RS conceived the project in discussion with MR. RS trained and supervised DAA, GMH, AA, ZK, LL, DP and ET, coordinated and designed experiments. RS and MR provided resources and funding. RS wrote the manuscript along with DAA, and all authors contributed to editing the final version of the manuscript.

## Data Availability Statement

The data that support the findings of this study are available from the corresponding author upon reasonable request.

## Supplementary Movie Captions

**Supplementary Movie 1.** An increase in GRABDA fluorescence is shown following bath application of 300 μM ACh in a live DLS brain slice from a mouse with Mito-GFP expressed in SNc astrocytic mitochondria. ACh is bath applied at 180 s to evoke dopamine release from SNc DA neuron axonal terminals in the DLS. Scale bar = 10 µm.

**Supplementary Movie 2.** An increase in GRABDA fluorescence is shown following bath application of 300 μM ACh in a live DLS brain slice from a mouse with Mito-GFP + Mito-PstI expressed in SNc astrocytic mitochondria. ACh is bath applied at 150 s to evoke dopamine release from SNc DA neuron axonal terminals in the DLS. Scale bar = 10 µm.

